# A multiscale pharmacokinetic-Boolean modelling framework for ibrutinib resistance in a Chronic Lymphocytic Leukaemia case study

**DOI:** 10.64898/2026.10.26.759125

**Authors:** Malvina Marku, Cécile Pineau, Marcelo Hurtado, Hugo Chenel, Jean-Baptiste Woillard, Loïc Ysebaert, Étienne Chatelut, Vera Pancaldi, Melanie White-Koning

## Abstract

Ibrutinib, a covalent Bruton’s tyrosine kinase (*BTK*) inhibitor, achieves durable remissions in chronic lymphocytic leukaemia (CLL), and yet a substantial proportion of patients relapse. A fundamental question is how the quantitative dynamics of drug exposure relate to the regulatory programmes and cell fate decisions that characterise response and relapse. Addressing it requires a multiscale framework that bridges the quantitative pharmacology of drug exposure with the molecular mechanisms encoded in the gene regulatory network, a coupling that has remained largely unexplored in the context of targeted therapy resistance. Here we present a multi-scale computational framework that addresses this question by coupling continuous patient-specific pharmacokinetic profiles with discrete Boolean dynamics, bridging two modelling paradigms that have so far operated in isolation. Applied to a longitudinal single-cell dataset from one CLL patient treated with ibrutinib across three clinical timepoints (pre-treatment, response, relapse), the framework integrates gene regulatory network inference, cell fate trajectory analysis, and a stochastic Boolean model, whose dynamics is driven by the patient-specific monthly drug exposure.

*AP-1* and integrated stress response regulons were specific to relapse, and the coupled model reproduced the clinical course. Its perturbation screens identified co-inhibition of *BTK* with *BCL2L1* or *NFKB1* as synergistic, in line with the rationale for dual targeting of *BTK* and the *BCL2* family in treatment-naive CLL. These results show how the framework turns single-patient data into mechanistic, testable hypotheses on resistance and combination therapy, which can be evaluated in larger cohorts and in experimental models.

## Introduction

Chronic Lymphocytic Leukaemia (CLL) is a malignancy of mature B cells characterised by the clonal expansion of CD19⁺CD5⁺ B lymphocytes that progressively accumulate in peripheral blood, bone marrow, and lymphoid tissues. Disease progression is driven by constitutive B-cell receptor (*BCR*) signalling, pro-survival interactions with the tumour microenvironment (1,2), and the acquisition of genetic alterations that confer growth and survival advantages (3). The introduction of ibrutinib, a first-in-class covalent inhibitor of Bruton’s Tyrosine Kinase (*BTK*), revolutionised the treatment of CLL by blocking *BCR*-driven survival signals, resulting in durable remission in a majority of patients (3). Despite this clinical success, a significant proportion of patients ultimately develop resistance to ibrutinib, limiting its long-term efficacy. Resistance to ibrutinib arises through two broadly distinct mechanisms. Acquired resistance is most commonly attributed to point mutations in *BTK* itself, in particular the *C481S* substitution that abolishes covalent drug binding, or to activating mutations in *PLCG2* that bypass BTK dependency (4,5). However, genetic mutations alone do not fully account for the clinical heterogeneity of resistance, and a growing body of evidence identifies non-mutational transcriptional plasticity as an equally important contributor. Leukemic cells can exploit compensatory signalling through the *TLR-MyD88*, *MAPK*, and *AP-1* pathways, reactivate lymph-node homing programmes centred on *CXCR4* and *RGS* proteins, and rewire the broader gene regulatory network (GRN) in ways that mitigate the effect of *BTK* inhibition (6,7). A fundamental question is how the quantitative dynamics of drug exposure relate to the regulatory programmes and cell fate decisions that characterise response and relapse. Addressing this question requires integrating information across multiple analytical layers: the structure of the gene regulatory network, the probabilistic fate trajectories that cells follow during treatment, and the quantitative drug exposure experienced by the malignant cells. Single-cell RNA sequencing (scRNA-seq) has opened a powerful window onto intraclonal heterogeneity and cell-state dynamics in CLL (8), but its integration with pharmacokinetic (PK) data, representing the actual drug concentration experienced by the malignant cells over time, remains largely unexplored. Because drug exposure varies substantially between patients (9), patient-level PK data provide a mechanistically grounded way to place longitudinal transcriptomic changes in the context of the exposure that each patient experienced. Recent computational approaches have begun to address this gap by inferring drug-responsive cell types from single-cell transcriptomic data through perturbation of inferred gene regulatory networks (10), and by reconstructing multiscale cell fate dynamics through the coupling of GRN-driven single-cell velocity with population dynamics (11), though integration with quantitative patient-specific pharmacokinetic profiles has not been previously achieved.

Mechanistic modelling adds a complementary layer to data-driven transcriptomics analysis. While GRN inference identifies the regulatory architecture under perturbations (e.g during treatment), it cannot fully encode the signalling logic that translates drug exposure into phenotypic outcome. Boolean models, which represent gene regulatory signalling interactions as discrete logical rules and simulate cell phenotypes as a probabilistic distribution over attractor states, serve as a powerful framework for this purpose in biology (12,13). A fundamental challenge in this field, however, is bridging the continuous quantitative scale of pharmacokinetics with the discrete logical rules of the Boolean model, which traditionally operates without a quantitative drug exposure dimension. Recent efforts in quantitative systems pharmacology (QSP) have begun to address this by embedding mechanistic signalling models with PK frameworks (14–17) and by integrating the cellular scales (18), but the integration of patient-specific PK profiles with single-cell transcriptomic data from patient samples within Boolean dynamic models, to our knowledge, has not previously been studied.

In our previous work (19), we reconstructed a context-specific GRN of CLL cells using longitudinal bulk RNA-seq data from patient-derived *in vitro* cultured Peripheral Blood Mononuclear Cells (PBMC), combining data-driven inference with curated transcription factor (TF)-target priors. The resulting network recapitulated key CLL pathways, including *BCR* and *NF-κB* signalling, and identified shared and patient-specific regulators governing temporal transcriptional responses. In another study (8), the authors conducted a longitudinal single-cell analysis on blood samples from a CLL patient undergoing treatment with the *BTK* inhibitor (ibrutinib). Using CITE-seq, which simultaneously profiles transcriptomes and surface proteins, they analysed PBMCs collected at three clinical time points: before therapy (M0), at early response after three months (M3), and at relapse after twenty-seven months of treatment (M27). The CITE-seq profiles revealed that the malignant CD19⁺CD5⁺ B-cell population is not homogeneous, but rather comprises at least three transcriptionally distinct leukaemic clusters, whereas at baseline, these clusters were present in different proportions. Following ibrutinib exposure, both leukemic B-cell and T-cell populations were substantially reduced, while NK cells’ abundance remained relatively stable. By the time of relapse at M27, the leukaemic population had re-expanded and was dominated by a clone harbouring a *BTK* mutation, which the authors detected in nearly all malignant cells. Pathway enrichment analysis demonstrated that, at relapse, the re-emergent clone exhibited strong upregulation of B-cell receptor signalling, *MYC*-driven programs, *NF-kB* signalling, and lymph-node homing pathways, all of which are in line with known mechanisms of ibrutinib resistance and aggressive disease behaviour (5,20,21).

Based on this longitudinal scRNA-seq dataset (8), we implement a multi-layered computational strategy combining (i) data-driven GRN inference to identify TFs with state-specific regulatory activity at each time point; (ii) trajectory inference to map cellular fate probabilities and identify genes and regulons associated with the stage transitions; (iii) a stochastic Boolean network model of CLL signalling, coupled to the patient’s ibrutinib PK profile and updated monthly, to mechanistically dissect the signalling dynamics underlying resistance; and (iv) a descriptive analysis relating gene expression and regulon activity to cumulative drug exposure (Fig. 1). Applied to a single patient as a case study, this framework identifies AP-1 and stress-response regulons as specific to relapse, reproduces the clinical course of treatment, and identifies the co-inhibition of *BTK* with *BCL2L1* or *NFKB1* as candidate combination strategies, illustrating how patient-level pharmacokinetics can be integrated with single-cell regulatory data and Boolean dynamics.

**Fig 1:**
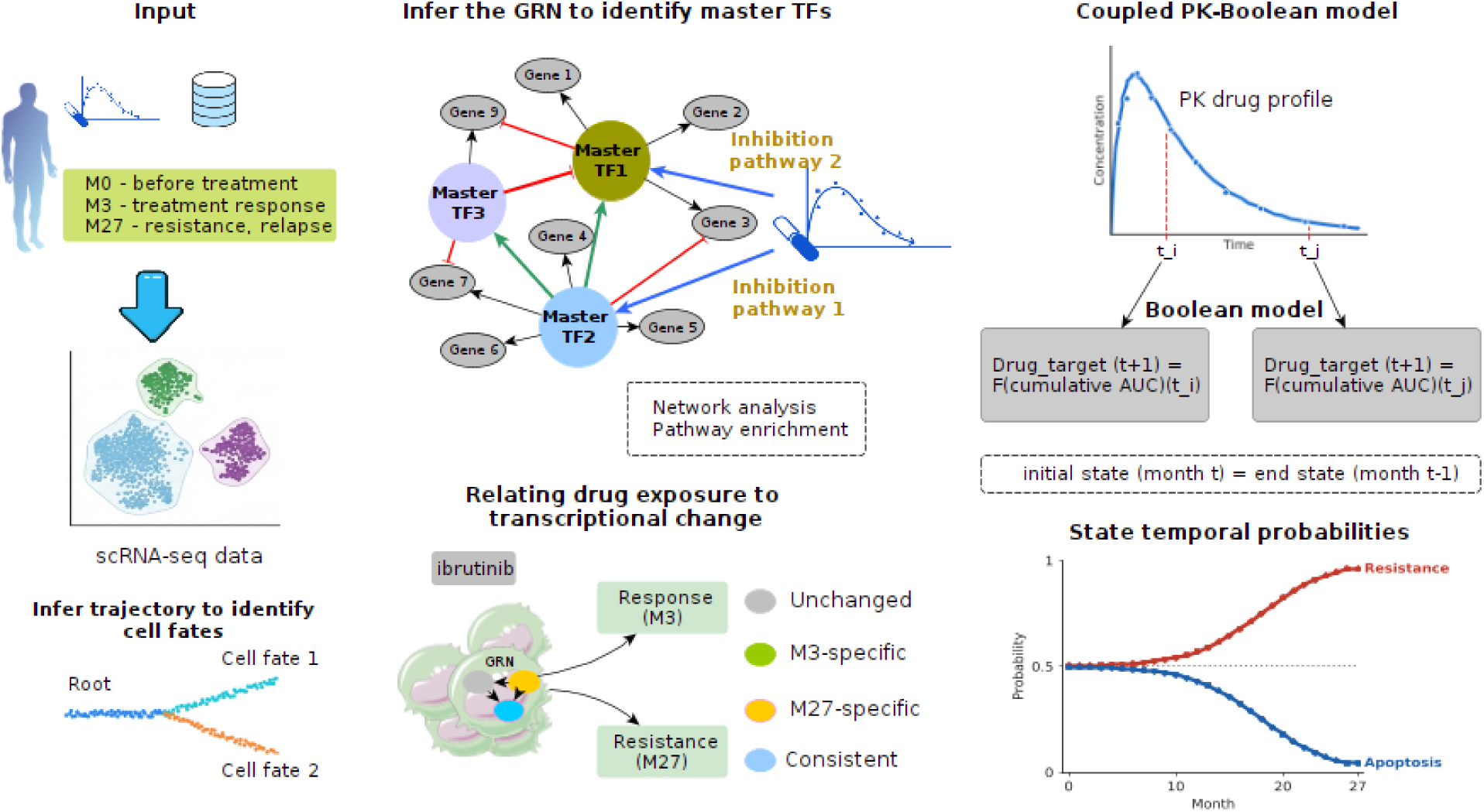
The main computational workflow. Longitudinal scRNA-seq data (M0, M3, M27) are used to infer the GRN and its master TFs and to map cell fates by trajectory inference. These are integrated with the patient’s ibrutinib PK profile and a coupled Boolean model to simulate phenotypic dynamics and screen for drug combinations.

## Results

Following the analysis performed on the longitudinal dataset in the original study, we focused on the B cell/CLL subpopulation. Non-coding genes and zero-variance genes were removed from the gene expression matrix, resulting in a matrix with 12872 genes and 10570 cells. As noted in the original publication, we observe a clear temporal separation of CLL cells across the three time points, with CLL cells from the M27 compartment more similar to those from the M0 compartment (Fig. S1).

### Data-driven GRN inference reveals putative markers of resistance

We employed pySCENIC (default values for all parameters and reference data (see Methods)) to identify the TFs that drive the expression of most of the genes in each time point, which are likely to play a particularly relevant biological role. The AUCell analysis revealed a total of 568 regulons with a significantly enriched motif for the corresponding TF. The regulon activity score for each cell type revealed a separate group of cells in relapse (Fig. 2 (a),(b)).

**Fig 2:**
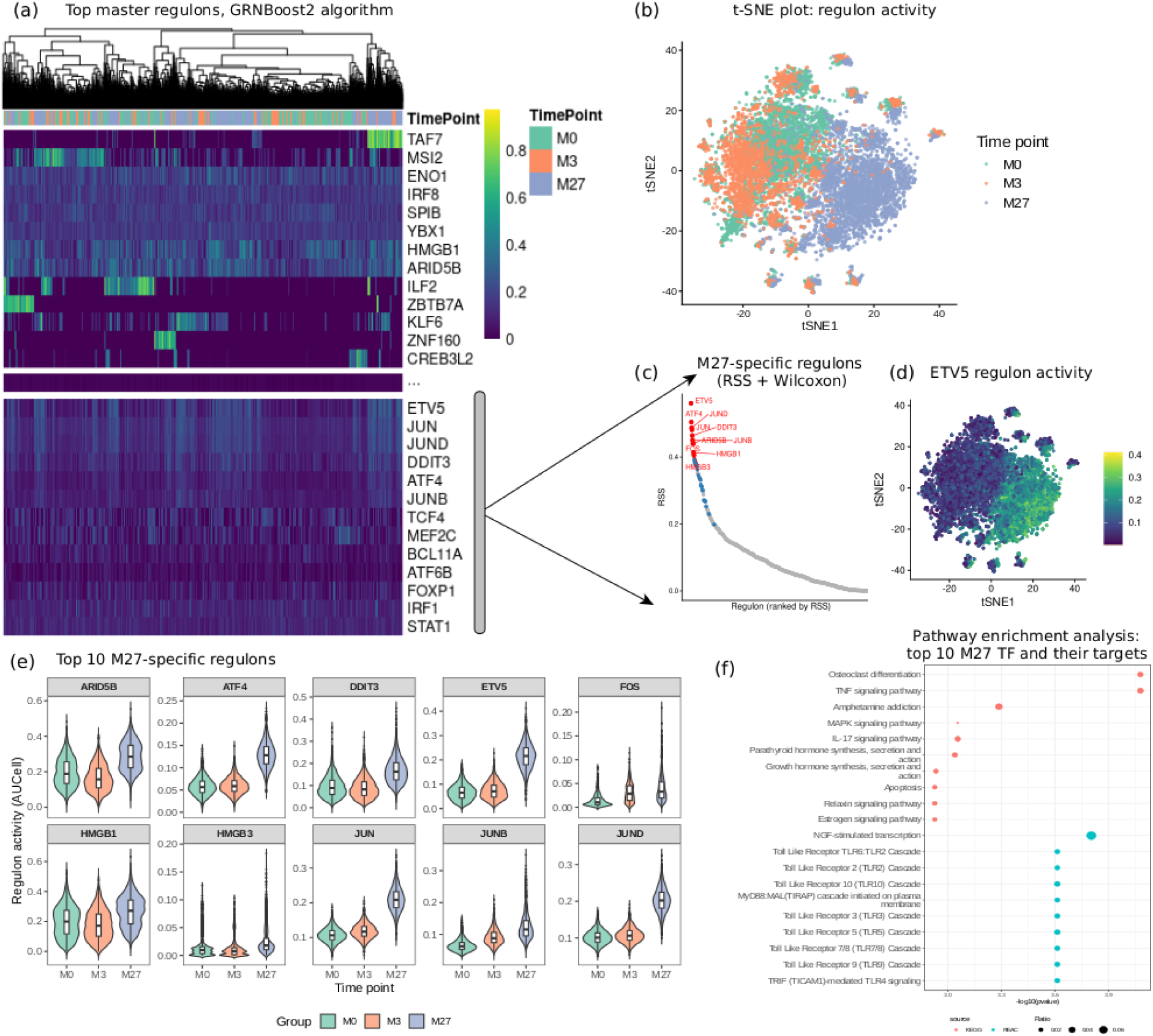
GRN inference identifies regulons that are specific to the relapse time point. **(a)** TF scoring based on the inferred GRN with pySCENIC. **(b)** t-distributed stochastic neighbour embedding (t-SNE) plot of CLL cells based on regulon activity from AUCell analysis, annotated by time point. **(c)** Top 10 M27-specific TFs based on regulon specificity score (RSS) and Wilcoxon test. **(d)** Regulon activity of *ETV5*. **(e)** Regulon score distribution of the top 10 M27-specific TFs in the three time compartments. **(f)** Pathway enrichment analysis on the top 10 M27-specific TFs and their targets.

We performed a Wilcoxon rank-sum test to identify the TF regulons up- or down-regulated in the M27 compartment, compared to M0 and M3. The analysis shows TFs such as *ETV5*, *ATF4*, *JUND*, *JUN*, *DDIT3*, *JUNB*, *FOS*, *ARID5B*, *HMGB1* and *HMGB3* among the top 10 with the highest regulon scores in M27 (*FDR* < 0. 05). In addition, we quantified the cell-type specificity of TF regulon scores by computing the Regulon Specificity Score (RSS) from the single-cell regulon activity matrix. For each cell group (M0, M3, M27), the distribution of regulon activities within that group is used to compute RSS values, which capture how selectively each regulon is activated within that group. Comparing the M27 compartment against M0 and M3 and ranking the regulons by their RSS, we identified key regulators driving transcriptional differences across the time points in our dataset (Fig. 2 (c)-(e), Fig. S2).

Performing pathway enrichment (GO and KEGG databases) of the top 10 up-regulated TFs in the M27 compartment and their targets, we observed a particular enrichment of *TLR* cascades (Fig. 2 (f)). These results suggest that these cells correspond to highly resistant cells, as it is often reported in the literature that the overexpression of *TLR* cascades in CLL is only partially inhibited by ibrutinib (6,22). In addition, the activation of *MyD88-MAL* adaptor complex, *MAPK* signalling and *IL-17* signalling reveal a strong over-representation of innate immune and stress-response signalling pathways. These results suggest that the M27 transcriptional program is driven by inflammatory and microenvironment-responsive signalling axes, rather than canonical B-cell receptor pathways alone. Activation of the *TLR-MyD88* signalling pathway is well-documented in CLL and has been associated with enhanced *NF-κB* and *MAPK* activation, increased survival, and resistance to apoptosis (7). The simultaneous enrichment of *MAPK* and *IL-17* signalling suggests amplification of pro-inflammatory transcriptional circuits that can promote proliferation, immune evasion, and adaptation to chronic stimulation (23).

Finally, examining the GRN structure (Supplementary file), we observe that TFs exhibiting high RSS in the M27 compartments also rank among the most central nodes in the inferred gene regulatory network, as assessed by degree and betweenness centrality (Fig. S3). This suggests that these factors not only display state-specific regulatory activity but also occupy key topological positions, coordinating information flow across multiple regulatory modules.

### Trajectory inference identifies M3 and M27 as separate cell fates

Following the workflow proposed in CellRank (24) (see Methods), we performed trajectory inference to calculate the macrostates and cell fate probabilities, setting the initial macrostate value to 5 (arbitrary). The analysis identified 3 major macrostates: one initial and two terminal states (Fig. 3 (a)), associating M0 cells with initial states and M3 and M27 states with terminal states, separating the responsive cells from the resistant ones (Fig. 3 (b), (c)). Employing Slingshot (25) and STREAM (26), we obtained similar results, with the two possible trajectories: M0 → M3 and M0 → M27 (Fig. S4).

**Fig 3:**
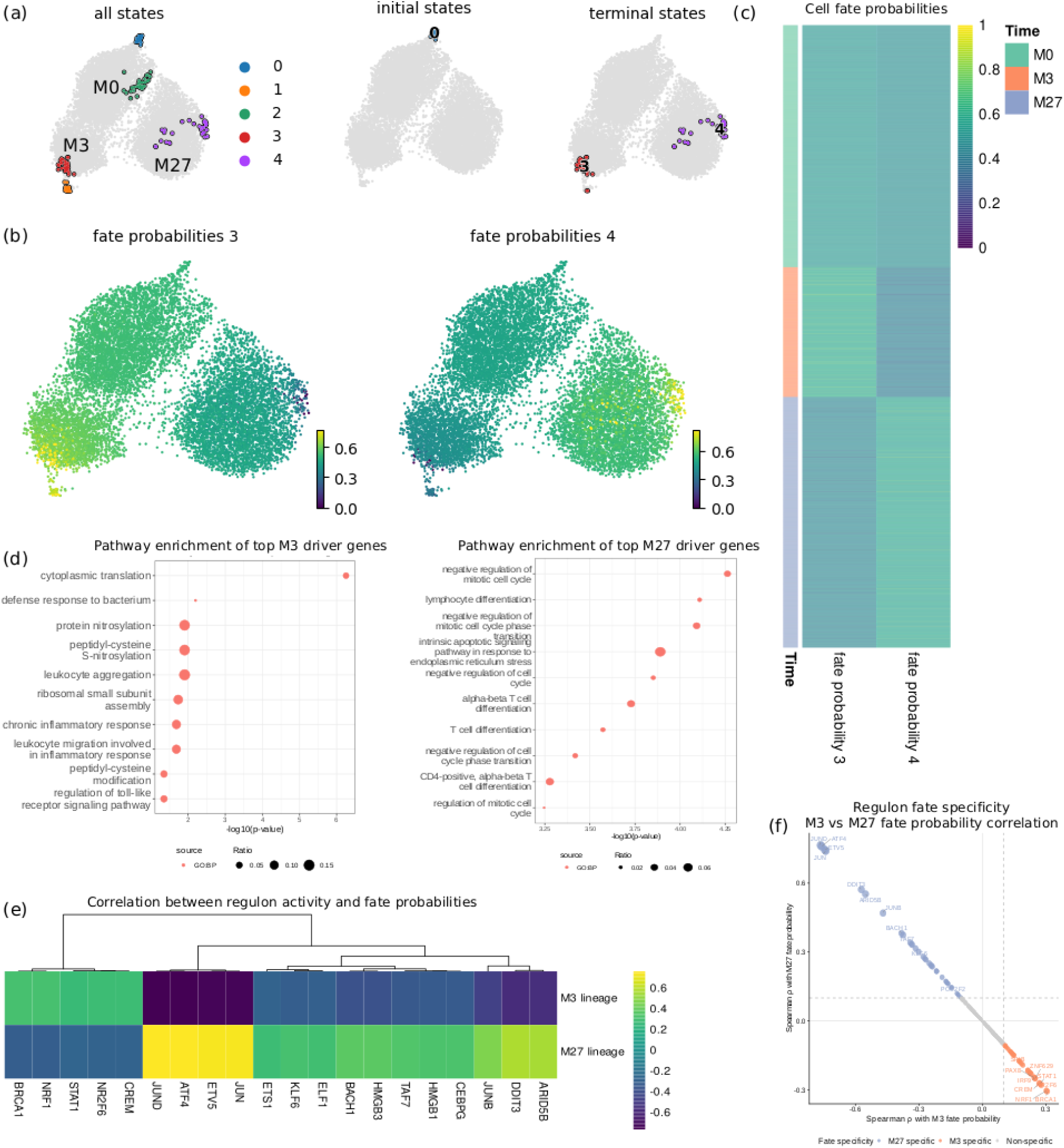
Trajectory inference separates the response (M3) and relapse (M27) fates and links them to distinct regulon programmes. **(a)** UMAP of the 5 macrostates and the three main macrostates identified: one initial and two terminals. **(b)** UMAP of terminal state probabilities. **(c)** Heatmap of cell fate probabilities highlighting the separation between M3 and M27 terminal states. **(d)** Pathway enrichment of top driver genes of M3 and M27 fates (GO database, BP ontology). **(e)** Correlation between cell fate probabilities and regulon activities. The heatmap shows the top 20 regulons for each fate. **(f)** Scatter plot of Spearman correlation between M3 and M27 fate probabilities. The TFs are classified as M3-specific, M27-specific or non-specific.

To identify fate-associated genes, we quantified the relationship between gene expression and fate probabilities across all cells. Specifically, for each terminal fate, we computed Spearman rank correlations between each gene’s expression level and the corresponding fate probability. We interpreted genes with high positive correlation coefficients as associated with progression toward that fate, whereas genes with negative correlation coefficients were associated with alternative trajectories.

The two fate-associated gene programmes were enriched for different biological processes, reflecting the divergent cellular fates of response and resistance. Genes driving the M0→M3 trajectory were primarily enriched for cytoplasmic translation and ribosomal small subunit assembly, alongside leukocyte aggregation and inflammatory response terms (Fig. 3 (d), Fig. S5 (a)). The translational enrichment reinforces a metabolically active, drug-responding cell population that sustains protein synthesis under ibrutinib pressure (27), while the inflammatory terms likely reflect sustained microenvironmental engagement at the early response phase.

Genes associated with the M0→M27 trajectory (Fig. S5 (b)), by contrast, were enriched for negative regulation of the mitotic cell cycle and cell cycle phase transition, and most notably for the intrinsic apoptotic signalling pathway in response to endoplasmic reticulum stress. The cell cycle arrest signature is consistent with CLL cells’ quiescence and drug-refractory state supported by *CXCR4*-mediated homing (20), a programme also identified during the GRN inference. The *ER* stress apoptotic term matches the integrated stress response (ISR) axis found by AUCell, where *ATF4* and *DDIT3*/*CHOP* were among the top M27-specific regulons: M27 fate-associated genes are both targets of ISR transcription factors and annotated to chronic *ER* stress adaptation (28).

Importantly, the top fate-associated genes of the M0→M27 trajectory included *RGS1*, *RGS2*, and *CXCR4*, previously reported as relapse markers in the original study (8), as well as *JUN* and *JUND*, independently identified as master regulators by the AUCell analysis. This convergence between data-driven trajectory inference and regulatory network analysis on the same set of genes provides additional support that these factors occupy a central position in the resistance transition.

### Linking fate probabilities to regulon activities identifies driver cell-fate regulons

To identify TFs whose regulatory activity is specifically associated with each terminal fate, we correlated AUCell values with fate probabilities across all cells. This integration allowed us to move beyond the static identification of M27-enriched regulons and ask which TFs are dynamically coupled to the probability of a cell committing to a given fate trajectory. To visualise the fate specificity of each regulon, we plotted the Spearman correlation with M3 fate probability against the correlation with M27 fate probability, positioning each TF in one of four quadrants: M27-specific, M3-specific, shared across both fates, or non-specific.

The analysis identifies largely non-overlapping sets of fate-associated regulons for the two terminal states, in agreement with the transcriptional divergence observed at the gene level. For the M3 response fate, the top correlated regulons included *BRCA1*, *NRF1*, *NR2F6* and *CREM*. Their functional enrichment (including oxidative stress response and mitochondrial biogenesis) is consistent with a drug-responsive cell population activating oxidative and inflammatory stress programmes, aligned with the translational and inflammatory enrichments observed in the M3 fate gene analysis. For the M27 resistance fate, the top correlated regulons recapitulate the findings of the AUCell analysis, with *AP-1* family members and integrated stress response TFs showing the strongest positive correlation with the probability of commitment to the resistance trajectory (Fig. 3 (d)-(f)).

### Boolean modelling of *BTK* signalling recapitulates CLL phenotypic dynamics and identifies synergistic therapeutic combinations

To mechanistically link patient-specific ibrutinib exposure to the phenotypic changes observed across the treatment course, and to identify nodes whose inhibition, alone or in combination, shifts cells toward apoptosis, we coupled the pharmacokinetic model to a stochastic Boolean network model of CLL signalling. The patient-specific PK data were incorporated in two main nodes: (i) BTK inhibition, encoded as a continuous transition rate derived from the pharmacokinetic model, and (ii) *PLCG2_mutant* probability, encoding acquired resistance mechanisms, modelled as a logistic function of the drug exposure (see Methods and Supplementary Material 1). The PK-Boolean coupled simulation (Fig. S6), initialised from AUCell-derived M0 transcriptomic states and run as a state-chained monthly sequence from M0 to M27, produces phenotypic attractor trajectories that are biologically interpretable and in line with the clinical treatment course (Fig. 4 (a)).

**Fig 4:**
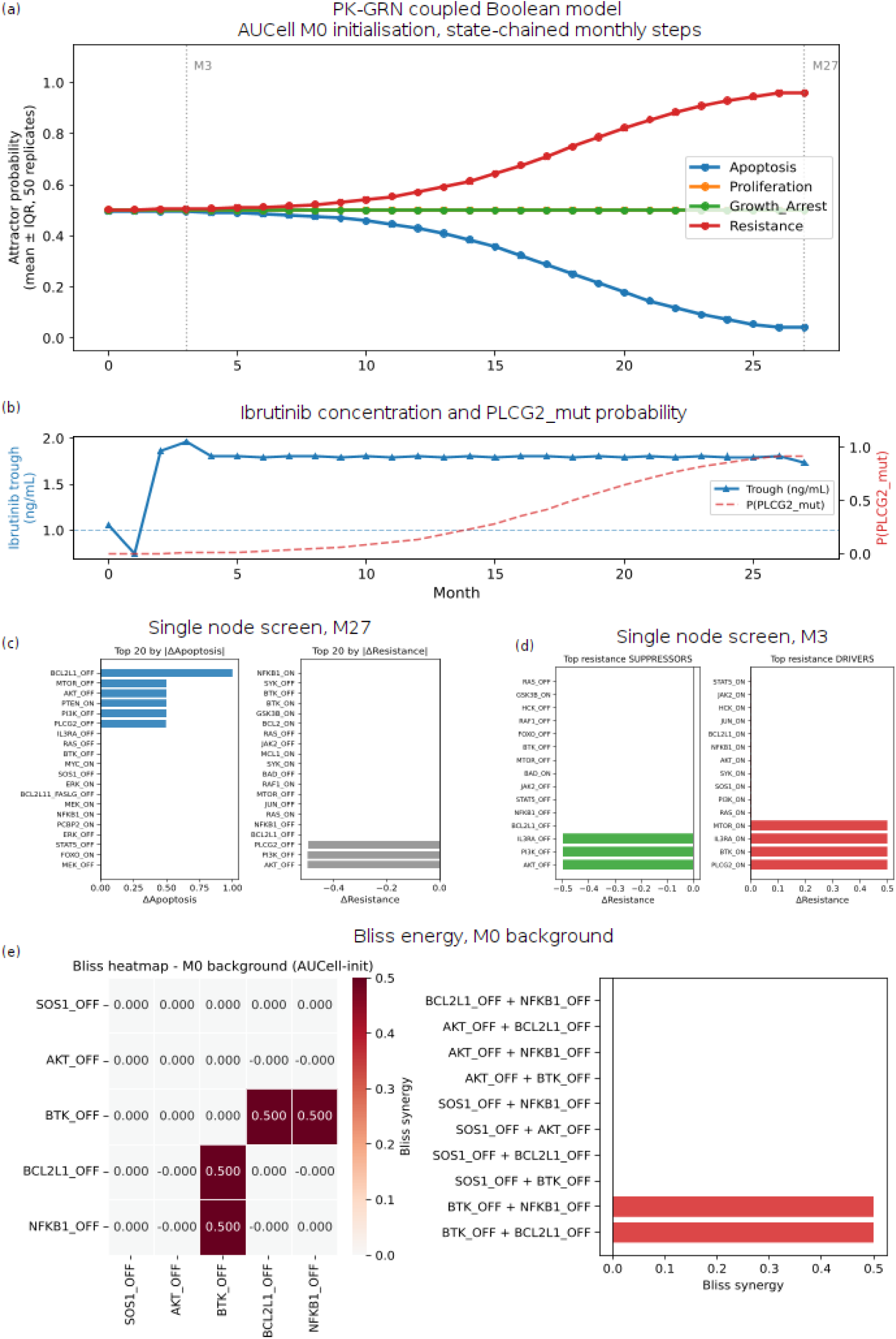
Boolean network modelling of CLL BCR/BTK signalling recapitulates phenotypic dynamics and identifies synergistic therapeutic targets. **(a)** Phenotypic attractor probability trajectories from M0 to M27, initialised from AUCell-derived M0 transcriptomic states. **(b)** Monthly ibrutinib concentration and derived P(*PLCG2_mut*) logistic expansion curve. **(c)** Single-node perturbation screen at M27: top 20 nodes ranked by ΔApoptosis (left) and ΔResistance (right). **(d)** Single-node perturbation screen at M3: top resistance suppressors (left) and drivers (right). **(e)** Bliss independence synergy screen at M0 background: heatmap (left) and ranked bar chart (right) showing *BTK_OFF* + *BCL2L1_OFF* and *BTK_OFF* + *NFKB1_OFF* as the top synergistic combinations.

At month 0, the model starts with a mixed CLL phenotype, with *Apoptosis* and *Resistance* both near 0.5, reflecting the neutral initialisation of unconstrained nodes and the competing pro-survival and pro-apoptotic signalling forces present in pre-treatment CLL cells. From month 2 onwards, as ibrutinib concentration remain approximately constant at 1.8 ng/mL (above the assumed IC50 of 1.0 ng/mL, giving ∼35% residual *BTK* activity) and the cumulative drug exposure increases monotonically, the *Apoptosis* probability begins a progressive decline while *Resistance* rises monotonically. By M27, *Resistance* reaches ∼0.95 and *Apoptosis* approaches 0, recapitulating the clinical picture of acquired resistance at relapse.

Importantly, the *BTK* inhibition rate derived from the PK profile remains essentially constant from month 2 to month 27 (Fig. 4 (b)), confirming that the phenotypic transition from the apoptotic to the resistant attractor is not driven by changing drug exposure. Instead, it is driven entirely by the logistic function of the *PLCG2_mutant* probability (P(*PLCG2_mut*)), which rises from near zero to ∼0.95 over the M12-M27 window as cumulative ibrutinib AUC crosses the AUC₅₀ threshold. This result shows that, in the model, the transition from the apoptotic to the resistant attractor is not activated by pharmacodynamic changes but by the progressive activation of the *PLCG2_mutant* signalling state, consistent with the emergence of the M27 resistance-associated transcriptional programme under constant drug pressure.

*Proliferation* and *Growth_Arrest* remain near 0.5 throughout the simulation, reflecting the bistable balance between the *MYC*/*JUN*-driven proliferative arm under partial *BTK* inhibition and the quiescent but viable phenotype characteristic of persisting CLL cells under ibrutinib treatment.

### Single-node perturbation screens

Single-node perturbation screens at M27 (Fig. 4 (c)) identified *BCL2L1_OFF* and *MTOR_OFF* as the strongest inducers of *Apoptosis* (ΔApoptosis≈0.9), followed by *AKT_OFF* and *PI3K_OFF*. This ranking reflects the central role of the *PI3K*-*AKT*-*mTOR*-*BCL2L1* survival axis in maintaining the M27 resistance attractor: when anti-apoptotic protection is removed by suppressing *BCL2L1* or *mTOR*, the network transitions almost entirely to the apoptotic basin. On the *Resistance* axis, *NFKB1_OFF*, *SYK_OFF*, and *BTK_ON* (partial reactivation) were the top suppressors, coherent with the *NF-κB* arm of the *Resistance* rule (*BCL2L1* and *NFKB1*) being the dominant contributor to *Resistance* probability at M27 once *PLCG2_mutant* is established.

At M3 (Fig. 4 (d)), *IL3RA_OFF* and *PI3K_OFF* emerged as the strongest resistance suppressors, reflecting the dependence of residual signalling on the *IL3→IL3RA→SYK→PI3K* axis when *BTK* is partially inhibited. The resistance driver screen identified *PLCG2_ON* and *IL3RA_ON* as the top drivers, both consistent with the model structure in which *IL3RA* and *PLCG2* are the two remaining active inputs to the *PI3K*-*AKT*-*NF-κB* cascade under ibrutinib. These results provide mechanistic validation that the model’s topology correctly captures the pharmacological vulnerabilities of CLL cells at each treatment stage.

### Bliss synergy screen identifies BTK + BCL2L1 co-inhibition as the top synergistic combination

Bliss scores were used to assess synergies between different types of *in silico* perturbations corresponding to combinations of inhibitors (see Methods). Starting from the M0 initial state, the pairwise Bliss independence synergy screen (Fig. 4 (e)) reveals that *BTK_OFF* + *BCL2L1_OFF* and *BTK_OFF* + *NFKB1_OFF* achieved the highest synergy scores (Bliss ≈ 0.5), substantially exceeding all other tested pairs. The Bliss heatmap confirms that these two combinations stand out clearly from the zero-synergy baseline, while all other pairs show additive or near-additive effects.

The *BTK_OFF* + *BCL2L1_OFF* result corresponds pharmacologically to ibrutinib combined with a *BCL*-*XL* inhibitor, a combination with documented preclinical activity in CLL and B-cell lymphoma (29). The *BTK_OFF* + *NFKB1_OFF* result similarly corresponds to ibrutinib combined with an *NF-κB* inhibitor, an axis of active therapeutic investigation in relapsed/refractory CLL.

Critically, this synergy is identified using M0 initial conditions, meaning that the model predicts that this combination would be most effective as an upfront strategy targeting the pre-existing resistance-associated transcriptional state before it has had the opportunity to expand under ibrutinib monotherapy. This recapitulates a broader pre-existing resistance process: if resistance is already present at diagnosis, then monotherapy sequential strategies are inherently limited, and co-targeting of the *BTK* and *BCL2L1*/*NF-κB* arms from the outset is mechanistically indicated.

### Integrating pharmacokinetic exposure with transcriptomics to dissect intrinsic and acquired drug resistance

To test whether ibrutinib exposure can account for the transcriptomic differences across the treatment course, we related patient-specific exposure to gene expression (focusing only on M3- and M27-fate driver genes) and TF regulon activity. Ibrutinib trough concentrations stabilised at about 1.8 ng/mL from month 2 and remained constant until M27 (Fig. 4 (b)). Drug concentration is therefore similar at M3 and M27, whereas the transcriptional state differs markedly between them (Fig. S1), so a change in concentration cannot explain the M3→M27 shift. Cumulative exposure, in contrast, increases with time on treatment. We therefore asked which genes and regulons change relative to their baseline level, and how they relate to cumulative exposure. We used two descriptive analyses: a regression-based classification of the temporal profile of genes and regulons, followed by their correlation with cumulative exposure (see Methods).

We first classified the 107 fate-driver genes by their deviation from the level predicted from M0 expression (Fig. 5 (a)). The M27-specific class contained *CXCR4*, *TSC22D3*, *CD69*, *PMAIP1*, *RGS2*, *KLF6* and *KLF2*, seven of the ten top M27 fate-associated genes (Fig. S5 (b)); *CXCR4* and *RGS2* are relapse markers of the original study (8). The M3-specific class included *IGLC2*, *BTG1*, *ZNF90* and *NHSL2*, of which *ZNF90* is among the top M3 fate-associated genes (Fig. S5 (a)).

**Fig 5:**
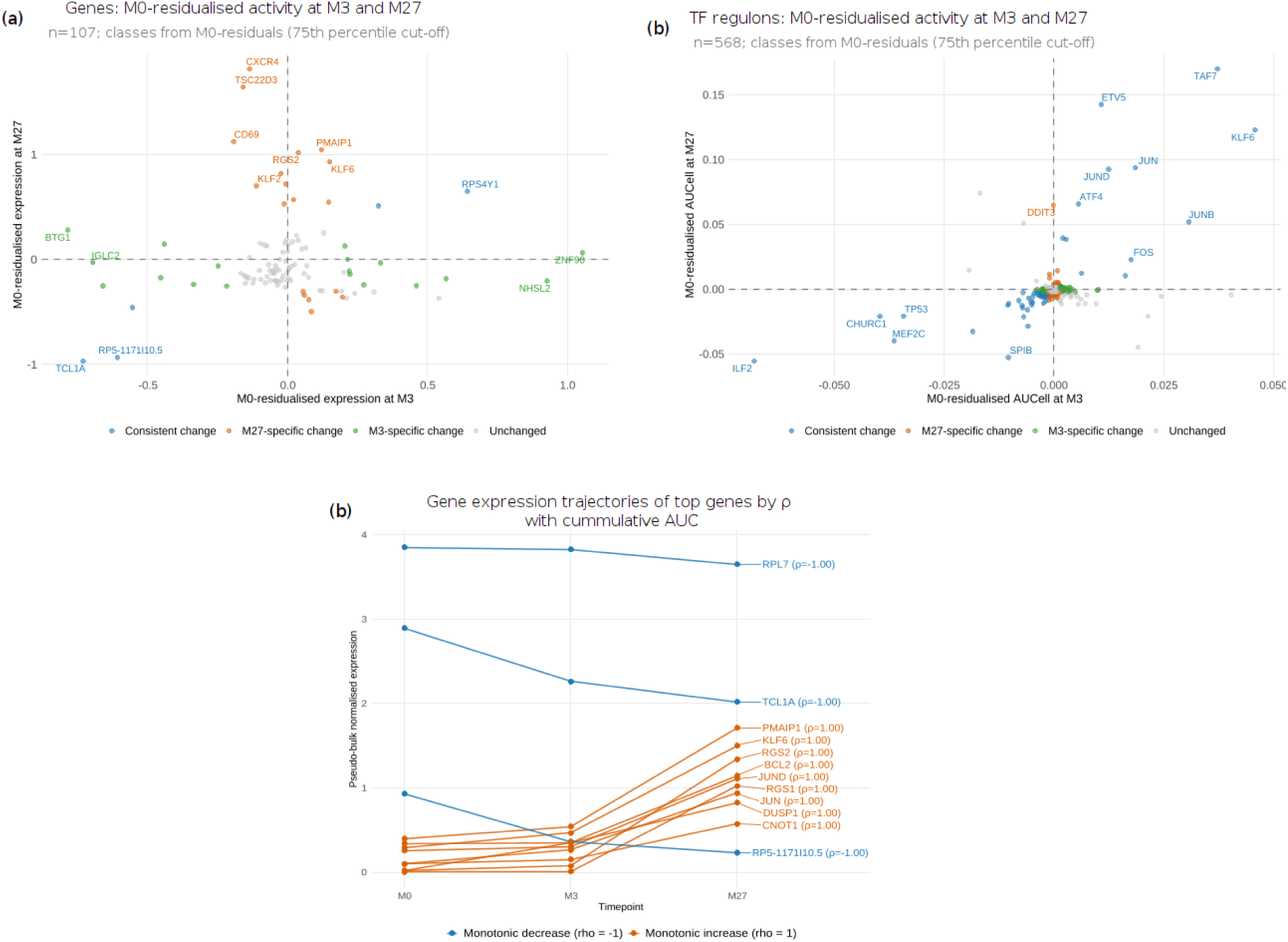
Relationship of gene expression and regulon activity to baseline and cumulative exposure. **(a)** Fate-driver genes (n = 107) classified by M0-residualised expression at M3 (x-axis) and M27 (y-axis): M27-specific change (orange), consistent change (blue), M3-specific change (green), unchanged (grey). **(b)** The same classification for TF regulons (n = 568), using AUCell scores. **(c)** Pseudo-bulk expression across M0, M3 and M27 of the genes with the strongest monotonic trends with cumulative AUC (ρ = ±1; orange, increase; blue, decrease).

The same classification of the 568 regulons (Fig. 5 (b)) showed that seven of the ten top M27-specific regulons from the GRN analysis (Fig. 2 (c),(e)) had positive residuals at both time points (consistent change): *JUN*, *JUND*, *JUNB*, *FOS*, *ATF4* and *ETV5*, together with *KLF6* and *TAF7*, whereas *DDIT3* showed M27-specific change. Regulons with lower than predicted activity at both time points included *TP53* and the B-cell transcription factors *SPIB* and *MEF2C*.

We then correlated each gene and regulon with cumulative AUC across M0, M3 and M27 (cumulative AUC = 0, AUC M0-M3, AUC M0-M27). With three time points, ρ can only take the values ±1 and ±0.5, and ρ = ±1 identifies features that change monotonically. Of the 107 genes, 30 (28.0%) increased, and 3 (2.8%) decreased monotonically; of the 568 regulons with non-zero activity at M0, 61 (11.0%) increased, and 109 (19.6%) decreased. The increasing genes and regulons (Fig. 5 (c)) largely overlapped with the M27-specific genes of panels (a) and (b). Decreasing genes included *TCL1A* and *RPL7*, which are among the top M3 fate-associated genes (Fig. S5 (a)). For most increasing genes, the change occurred between M3 and M27, with little change between M0 and M3.

The genes and regulons that characterise the relapse state in the GRN and trajectory analyses (Fig. 2, 3 and S5) therefore also increase monotonically with cumulative exposure. However, because cumulative AUC is proportional to time on treatment after month 2, these associations cannot distinguish drug-induced adaptation, selection of cells already present, and disease progression or sample-specific effects. Taken together, although drug concentration cannot explain the transcriptional shift between M3 and M27, the two analyses describe which genes and regulons change during treatment, indicating a potential association with treatment duration.

## Discussion

The central question motivating this study was whether ibrutinib resistance in CLL reflects a transcriptional programme that is pre-encoded in the heterogeneous pre-treatment cell state or one that emerges progressively under pharmacological pressure. By integrating single-cell gene regulatory network inference, trajectory analysis, Boolean network modelling, and patient-specific pharmacokinetic data across a longitudinal CITE-seq dataset, we obtain evidence that resistance in this patient is predominantly intrinsically determined. Three observations support this interpretation: M27 cells do not originate from the M3 drug-response state in the trajectory inference; ibrutinib exposure, which plateaus from month 2, does not explain the transcriptional shift between M3 and M27, whereas the Boolean network model, initialised from pre-treatment initial conditions and driven by constant drug exposure, reproduces the progression to the resistance attractor. These results come from methodologically distinct analyses, although the GRN and trajectory analyses derive from the same single-cell dataset, and all are limited to a single patient.

### Coupling pharmacokinetic exposure with transcriptomic dynamics reveals a flat-exposure resistance mechanism

A key finding of the PK-transcriptomic integration is that ibrutinib plasma concentrations, despite spanning a clinically relevant range in the first months of treatment, do not explain transcriptomic variation between M3 and M27. The regression analysis shows strongly negative ΔR² values for all prediction tasks, indicating that adding PK features to baseline expression actually reduces predictive performance: a result that is counterintuitive at first but becomes interpretable in light of the flat PK profile. Ibrutinib trough concentrations stabilise at approximately 1.8 ng/mL from month 2 onwards and show minimal variation through M27, providing no pharmacokinetic variance to explain the transcriptomic shift. This result is consistent with the known pharmacology of ibrutinib: as a covalent inhibitor, ibrutinib achieves near-complete BTK occupancy at *C_max_* even at standard doses, and the steady-state trough reflects a pharmacological plateau rather than a dynamic process. The M3→M27 transcriptomic shift is therefore not a pharmacodynamic response to changing drug levels, but a consequence of a resistance-associated transcriptional shift operating under constant pharmacological pressure: precisely the scenario that the logistic *PLCG2_mutant* expansion model in the Boolean simulation is designed to represent.

This finding has a methodological implication beyond this specific patient. The assumption that PK variability should predict transcriptomic variability in covalent inhibitor therapy may be incorrect in patients who have achieved steady-state drug exposure, as the relevant driver of transcriptional change shifts from pharmacodynamics to clonal dynamics. PK-transcriptomic integration frameworks should therefore incorporate explicit models of the timing of resistance-state transitions (as done here through the *PLCG2_mutant* logistic model) rather than treating drug exposure as the sole explanatory variable for resistance-associated transcriptional change. While recent approaches have demonstrated the feasibility of inferring drug-responsive cell types from untreated scRNA-seq data via GRN perturbation (10), the present work extends this paradigm by grounding the drug effect in quantitative patient-specific pharmacokinetic exposure rather than binary perturbation assumptions.

### Therapeutic implications of the Boolean network predictions

The Bliss synergy analysis from M0 initial condition identifies *BTK_OFF* + *BCL2L1_OFF* as the most synergistic combination (Bliss≈0.5), corresponding pharmacologically to ibrutinib combined with a *BCL*-*XL* inhibitor such as navitoclax or with venetoclax targeting *BCL2*. This prediction is in agreement with preclinical evidence for the ibrutinib-venetoclax combination in CLL and B-cell lymphoma (30) and with emerging clinical data from the GLOW (31) and CAPTIVATE (32) trials, which demonstrated superior outcomes for ibrutinib + venetoclax doublet therapy compared to ibrutinib monotherapy in treatment-naive CLL (33,34). Critically, our model generates this prediction from the M0 background, meaning it identifies this combination as most effective as an upfront strategy targeting the pre-existing resistant clone before it expands under monotherapy. This is mechanistically coherent with a *BCR*-independent, *JUN*/*AP-1*/*BCL2L1*-active transcriptional state already detectable at M0 (above): monotherapy ibrutinib would be expected to favour cells already expressing this survival programme, regardless of how complete *BTK* inhibition is, because the relevant survival signal has already bypassed *BTK*. Co-targeting *BCL2L1* from the outset would, in principle, remove the anti-apoptotic protection that this resistance-associated state relies on, potentially contributing to the prevention of resistance emergence, a hypothesis that requires experimental validation in patient-derived models before clinical translation.

The single-node perturbation screen at M27 provides a complementary therapeutic perspective. *BCL2L1_OFF* and *MTOR_OFF* are the strongest apoptosis inducers at the resistance stage, reflecting the centralisation of survival signalling around the *PI3K-AKT-mTOR-BCL2L1* axis once *PLCG2_mutant* is established. This identifies *mTOR* inhibition as a potential salvage strategy at relapse, corroborating clinical case evidence for *mTOR* pathway activation in ibrutinib-resistant CLL (35) and with ongoing clinical evaluation of *PI3K*-*mTOR* inhibitors in relapsed/refractory disease (36).

### Limitations

#### Several limitations of this study should be acknowledged

<u>First</u>, the analysis is based on a single patient, which precludes generalisation and means that all findings are mechanistically plausible but not statistically validated across a population. The intrinsic-resistance interpretation cannot be confirmed without genomic or single-cell DNA sequencing data to determine whether cells expressing this resistance-associated transcriptional programme carry *BTK C481S* or *PLCG2* mutations, or whether the phenotype is purely transcriptional (non-genetic). Furthermore, while our results point to a predominantly pre-existing resistance programme, we cannot fully exclude the contribution of drug-induced transcriptional adaptation occurring in parallel with the observed transcriptional shift. Recent work has demonstrated that targeted therapy can itself trigger sequential *NF-κB*-driven transcriptional waves that promote early drug tolerance before genetic resistance emerges (37), a mechanism that may operate alongside the pre-existing transcriptional programme we identify here and that cannot be disentangled from it in the absence of dense longitudinal sampling within each treatment phase.

<u>Second</u>, additional limitations are inherent to the Boolean network modelling framework. The *PLCG2_mutant* logistic expansion model additionally contains two free parameters (AUC₅₀ fraction and sigmoid steepness) that are calibrated manually to the observed clinical progression timepoint rather than fitted to data; a population-level analysis with multiple patients would enable proper parameter estimation and testing of whether cumulative AUC is the relevant mechanistic driver of clonal expansion. In addition, the Bliss synergy *in silico* predictions have not been experimentally validated in patient-derived CLL cells and should be interpreted as hypotheses for experimental testing rather than clinical recommendations.

<u>Third</u>, the analysis relating gene expression to drug exposure is descriptive and limited by the design of the study. Ibrutinib trough concentrations stabilised at approximately 1.8 ng/mL from month 2 and remained essentially constant until M27 (Fig. 4 (b)), so the M3→M27 shift cannot be explained by a change in concentration. This plateau, however, is a model prediction: concentrations were measured only during the first months of treatment and were simulated from the population PK model afterwards. Cumulative exposure, in contrast, increases with time on treatment, and because all time points after M0 are on treatment, it cannot be separated from treatment duration or from other processes that change over time, such as selection of cells already present, disease progression, and differences between the three samples. With three time points, the Spearman correlation can only take the values ±1 and ±0.5 and cannot be tested for significance, so genes with ρ = ±1 are those that change monotonically over time and not genes shown to respond to exposure. The gene set was also selected for association with the M3 or M27 fate, so an increase of these genes over time is expected. Identifying genes whose expression depends on exposure would require exposure to vary independently of time, for example in a multi-patient cohort or a dose-response experiment.

## Conclusion

We present a multi-scale computational framework integrating single-cell gene regulatory network inference, trajectory analysis, stochastic Boolean network modelling, and patient-specific pharmacokinetic data to dissect the mechanisms of ibrutinib resistance in CLL. Applied to a longitudinal CITE-seq dataset spanning pre-treatment, early response, and relapse, the framework reveals that resistance in this patient is predominantly intrinsically determined and detectable ahead of relapse in the trajectory analysis and is reproduced by a mechanistic Boolean model initialised from pre-treatment transcriptomic states, without requiring drug-induced pharmacodynamic adaptation to explain its emergence.

The coupled PK-Boolean model recapitulates the phenotypic dynamics of the treatment course and identifies *BTK + BCL2L1* co-inhibition as the most synergistic therapeutic combination from the pre-treatment cell state. This prediction is in agreement with the clinical benefit observed for upfront ibrutinib-venetoclax doublet therapy and provides a mechanistic rationale for co-targeting the BCR bypass and anti-apoptotic arms simultaneously, rather than sequentially, in patients whose pre-treatment tumour already expresses a *BCL2L1*-dependent resistance-associated transcriptional programme.

Ultimately, the computational framework developed here is designed to be applicable to other targeted therapies and cancer types, where the distinction between pre-existing and acquired resistance has direct therapeutic implications, and its generalisability should be assessed in future multi-patient studies where longitudinal single-cell data and patient-level drug exposure data are simultaneously available. Future work will extend this framework to patient cohorts to test whether the pre-existing resistance signature identified here generalises across CLL patients, and to validate the predicted combination synergies in patient-derived cell models.

## Methods

### Gene regulatory network (GRN) inference with pySCENIC

We implemented pySCENIC (38,39) to infer the GRN of the CLL cells from single-cell data on all cells pooled across the three time points. In principle, pySCENIC uses a gene expression dataset and a list of TFs and employs the GRNBoost2 (40) or GENIE3 (41) algorithms to infer the causal regulatory relationships between TFs and their targets. Furthermore, a critical step of the pySCENIC workflow is the pruning of the false positive interactions, using cis-regulatory regions, implemented in the RcisTarget module with default parameters (NES > 3.0, auc_threshold = 0.01). Lastly, based on the inferred and pruned network, the workflow incorporates an estimation of TF scores via the AUCell module, which estimates, for each cell, the enrichment of a regulon’s target genes among the most highly expressed genes in that cell, expressed as an area-under-the-recovery-curve score, yielding a continuous, per-cell activity score for each inferred regulon.

In our case, we employed the standard pySCENIC workflow, using the GRNBoost2 algorithm for GRN inference with default parameters.

The reference files for performing GRN inference with pySCENIC were downloaded from https://resources.aertslab.org/cistarget/:

• Databases: [hg19-500bp-upstream-7species.mc9nr.feather](https://resources.aertslab.org/cistarget/databases/old/homo_sapiens/hg19/refseq_r45/mc9nr/gene_based/), [hg19-tss-centered-5kb-7species.mc9nr.feather](https://resources.aertslab.org/cistarget/databases/old/homo_sapiens/hg19/refseq_r45/mc9nr/gene_based/), [motifs-v9-nr.hgnc-m0.001-o0.0.tbl](https://resources.aertslab.org/cistarget/motif2tf/)

• TF list: https://resources.aertslab.org/cistarget/tf_lists/allTFs_hg38.txt

To identify TFs whose regulatory activity is preferentially concentrated within a specific timepoint or group of cells, we computed the Regulon Specificity Score (RSS) for each regulon (42). Briefly, RSS quantifies how selectively a regulon’s activity is confined to a given cell group by comparing the regulon’s normalised per-cell activity distribution (from AUCell) against an idealised distribution restricted entirely to that group, using the Jensen-Shannon divergence between the two (*RSS* ∈ [0, 1], higher scores indicate greater group-specific activity). RSS*_score_* was computed separately for each timepoint (M0, M3, M27) from the single-cell regulon activity matrix.

#### Trajectory inference with CellRank

Trajectory inference and cell fate bias estimation were performed using CellRank (24), a graph-based probabilistic framework for inferring cellular state transitions from single-cell RNA-sequencing data. As spliced and unspliced transcript counts were not available, analyses were conducted using the *ConnectivityKernel* CellRank class, which leverages transcriptional similarity encoded in the k-nearest neighbour (kNN) graph without relying on RNA velocity information.

Briefly, gene expression was normalised (log-normalised, SCTransform), and the top 2000 most variable genes were identified and used to reduce dimensionality via principal component analysis (PCA, *n_PC_* = 30, default). To model state transitions between cells, we used the *ConnectivityKernel* from the kNN graph (*n _neig_*_ℎ*bours*_ = 30, default), which defines a Markov transition matrix in which transition probabilities are proportional to transcriptional similarity between neighbouring cells. In the absence of RNA velocity, this approach captures the intrinsic geometry of the cellular manifold, enabling inference of relative progression and state stability from the graph topology. Cellular macrostates and fate structure were inferred using Generalised Perron Cluster Analysis (GPCA) applied to the transition matrix. Within this framework, *initial states* were defined as cells with low incoming and high outgoing transition probabilities, *terminal states* were defined as absorbing or near-absorbing states in the Markov process, corresponding to stable transcriptional endpoints, and remaining macrostates were intermediate. This graph-based definition does not assume explicit temporal ordering but instead reflects probabilistic flow on the cellular neighbourhood graph.

For each cell, fate probabilities toward each identified terminal macrostate were computed. These probabilities quantify the likelihood that a random walk starting from a given cell will ultimately reach a particular terminal state, providing a soft assignment of cells to alternative differentiation outcomes. Fate probabilities were visualised on the Uniform Manifold Approximation and Projection (UMAP) embedding to reveal continuous trajectories and branching structures across the cellular landscape. Because RNA velocity information was not available, inferred trajectories represent state hierarchies and fate biases rather than explicit temporal dynamics. Consequently, we interpret the results in terms of transcriptional commitment, stability, and relative progression along differentiation, rather than absolute developmental time.

Slingshot (25) and STREAM (26) were additionally run as alternative methods to validate the CellRank trajectories.

### Pharmacokinetic (PK) model of ibrutinib

In order to integrate drug kinetics into CLL cells’ phenotypic dynamics during treatment, we used a previously published population-PK model of ibrutinib and its metabolite, Dihydrodiol-Ibrutinib (DHD-ibrutinib) (43). In this study, plasma concentrations from 89 patients treated for lymphoid malignancies, including the patient for whom the CITE-seq data were generated, were collected over 6 months and analysed using nonlinear mixed-effects modelling. The PK of ibrutinib and its metabolite were fitted using a two-compartment model with sequential zero-order then first-order absorption (after a lag time), also capturing first-pass hepatic metabolism and the complex disposition dynamics.

Building on this model, patient-specific ibrutinib concentration-time profiles were simulated for the full treatment course. Cumulative drug exposure was quantified as the area under the concentration-time curve (AUC), computed using the linear trapezoidal rule:

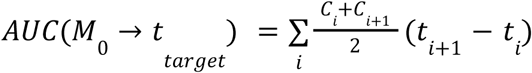

where *C^i^* and *C_i_*_+1_ are consecutive predicted plasma concentrations at *t_i_* and *t_i_*_+1_ time points. Three cumulative exposure metrics were derived by setting the upper integration limit *t_target_* to month 3 or month 27: *AUC_M_*_3_ (M0 to M3), capturing early treatment exposure; *AUC_M_*_27_ (M0 to M27), capturing total lifetime exposure; and *AUC_late_* (M3 to M27), capturing late-phase exposure during the resistance window.

### Boolean network model construction

A Boolean network model of CLL signalling under ibrutinib treatment was constructed by adapting the Bio Model Analyser leukaemia signalling network (44). The complete steps for building the model are described in detail in Supplementary Material 1. The model was implemented in MaBoSS (45) and executed within the CoLoMoTo Docker computational environment (46). MaBoSS performs stochastic simulation of Boolean networks, sampling trajectories under continuous-time Markov dynamics governed by node-specific up/down transition rates, and reporting the resulting probability distribution over attractor states. In our case, four phenotypic output nodes were defined: *Apoptosis*, *Proliferation*, *Growth_Arrest*, and *Resistance*. The final network contains 37 nodes (Fig. S.6), whereas the full details on model building steps can be found in Supplementary Material 1.

Three simulation conditions were defined to represent the CLL ibrutinib treatment time course: M0, M3, and M27. Two categories of pharmacological constraints were applied as condition-specific locks:

i. *BTK* inhibition, encoded as a continuous transition rate derived from the pharmacokinetic model (rather than a binary lock), reflecting the incomplete occupancy achieved at therapeutic trough concentrations (full description of the model parameters can be found in Supplementary Material 1):

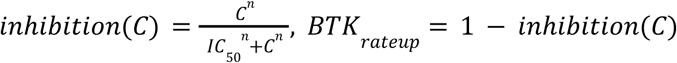

with *IC*_50_ = 1. 0 *ng*/*mL* and Hill coefficient *n* = 1, since ibrutinib is a covalent inhibitor where the kinetic *IC*_50_ for irreversible bond formation approximates a first-order occupancy model (47,48).

(ii) condition-specific biological states, including *PLCG2_mutant*=ON and *BCL2*/*MCL1*=ON at M27 to encode acquired resistance mechanisms (calibrated so that clonal dominance coincides with the observed clinical resistance timepoint):

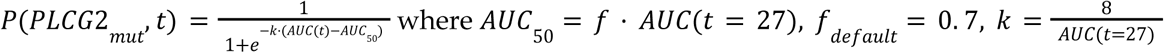

To ground the Boolean model’s initial states in single-cell transcriptomic data, regulon enrichment scores derived from AUCell analysis were used to initialise TF node states; the remaining nodes were not directly constrained by transcriptomic data at initialisation and were put at random states (*prob*([*node*] = 1) = *prob*([*node*] = 0) = 0. 5). The AUCell scores were binarised per regulon using an automatically selected threshold (valley of bimodal distribution if Sarle’s bimodality coefficient BC > 0.555, otherwise discriminability-maximising threshold), and then aggregated to a majority-vote consensus per timepoint.

Three main simulations were run under condition-specific initial states (M0, M3, M27) to characterise the phenotypic attractor for each treatment stage. Attractor probabilities for all four output nodes were extracted at the final time point and used as reference values for all subsequent perturbation analyses. We additionally performed single-node perturbation screens (systematically fixing each node ON or OFF and quantifying the resulting change in phenotypic attractor probabilities) at M3 and M27, and a pairwise Bliss independence synergy screen across candidate target-node combinations from the M0 background. Full parameter values, the complete list of perturbed nodes, and the methodology for each simulation and knockout are provided in Supplementary Material 1.

### Relating pharmacokinetic exposure to transcriptomic change

Cumulative ibrutinib exposure was computed as the area under the simulated concentration-time curve (AUC, trapezoidal rule) from M0 to M3 and from M0 to M27. For each gene and each regulon, pseudo-bulk values were obtained by averaging normalised expression (genes) or AUCell regulon scores (regulons) across all cells of each time point. The analysis was restricted to fate-driver genes (those with a correlation above 0.1 with the M3 or M27 fate probability in the trajectory analysis) and to regulons with a non-zero mean activity at M0. To describe the temporal profile of each feature, M3 and M27 values were regressed on M0 values across genes (or regulons), and each feature was classified from its residuals at the two time points (75th percentile of the absolute residuals as the cut-off). Features with a large M27 residual and a small M3 residual were labelled M27-specific, those with large residuals of the same sign at both time points were labelled as showing consistent change, and those with a large M3 residual and a small M27 residual were labelled M3-specific; all other features were considered unchanged.

To relate expression to exposure, the Spearman correlation was computed between the three pseudo-bulk values of each feature (M0, M3, M27) and cumulative AUC (0, AUC(M0→M3), AUC(M0→M27)). Because only three time points are available, ρ can only take a few discrete values, and ρ = ±1 identifies features that change monotonically over the treatment course. As cumulative AUC increases monotonically with time on treatment, this analysis is descriptive as it cannot separate exposure from treatment duration.

## Data and code availability

The R scripts and Jupyter notebooks used for performing all the analysis and reproducing the figures are available at https://github.com/M-Marku/CLL_GRN_sc_multiscale.git. Libraries and environment versions can be found in Supplementary Material 2. Large output files (single cell objects and count matrices) are available at the Zenodo repository https://doi.org/10.5281/zenodo.23320593.

The original processed CITE-seq data can be found in the original publication (8).

## Acknowledgments

The authors thank the Genotoul Bioinformatics computation cluster (https://bioinfo.genotoul.fr/) for enabling access to computational resources for performing various analyses. The authors thank Loïc Paulevé, Laurence Calzone, Augustine Blanc-Boekholt and Victoria Bruning for their constructive discussions.

## Author contributions

Conceptualisation: M.M; Computational workflow: M.M; Data exploration: M.M, M.H, C.P, H.C; Project supervision: M.W-K, E.C; Interpretation of results: M.M, C.P, L.Y, V.P, J-B.W, M.W-K; Writing original draft: M.M; Review and editing: all authors.

## Supplementary Figures

**Figure S1:**
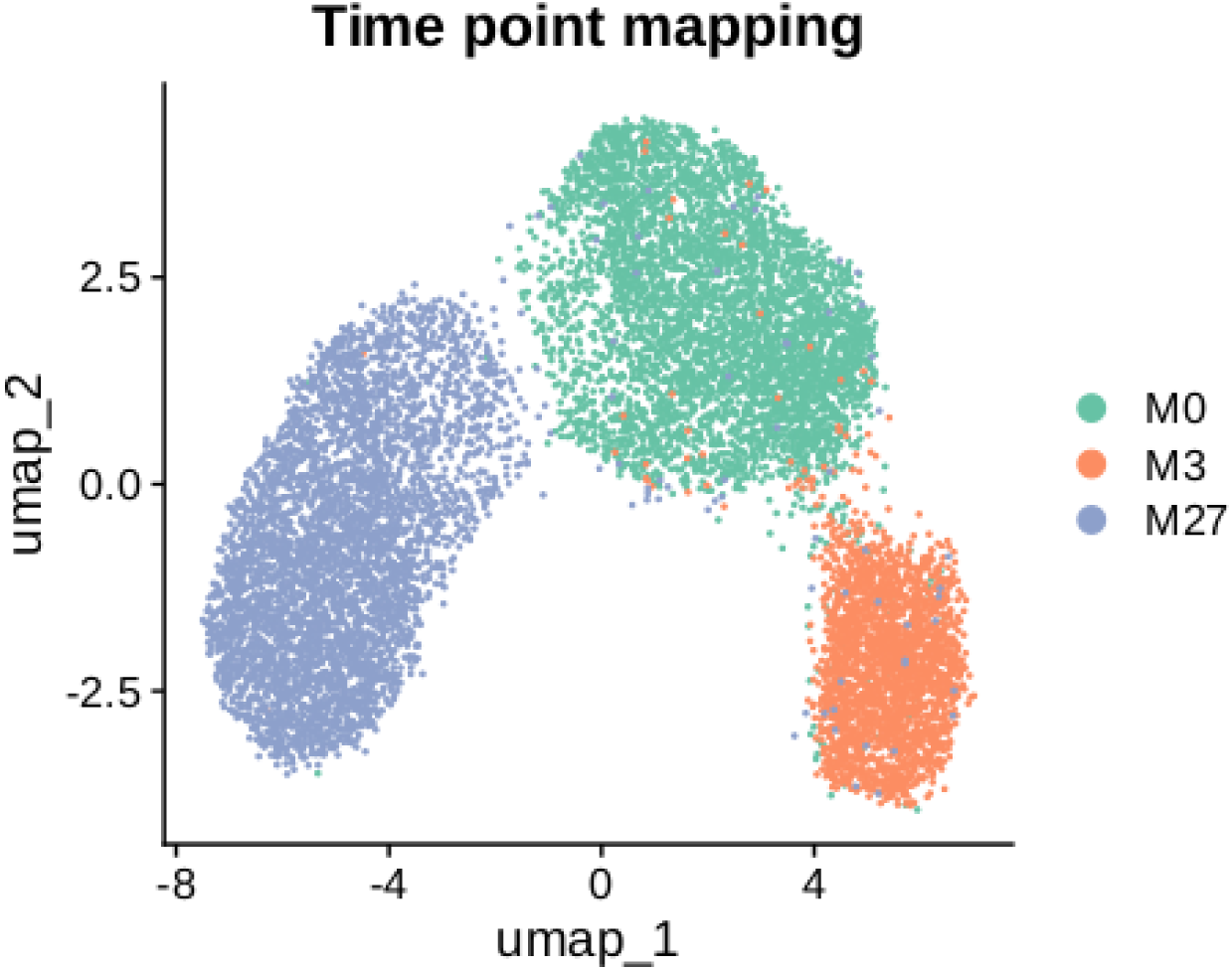
Time point mapping on the CLL subpopulation.

**Figure S2:**
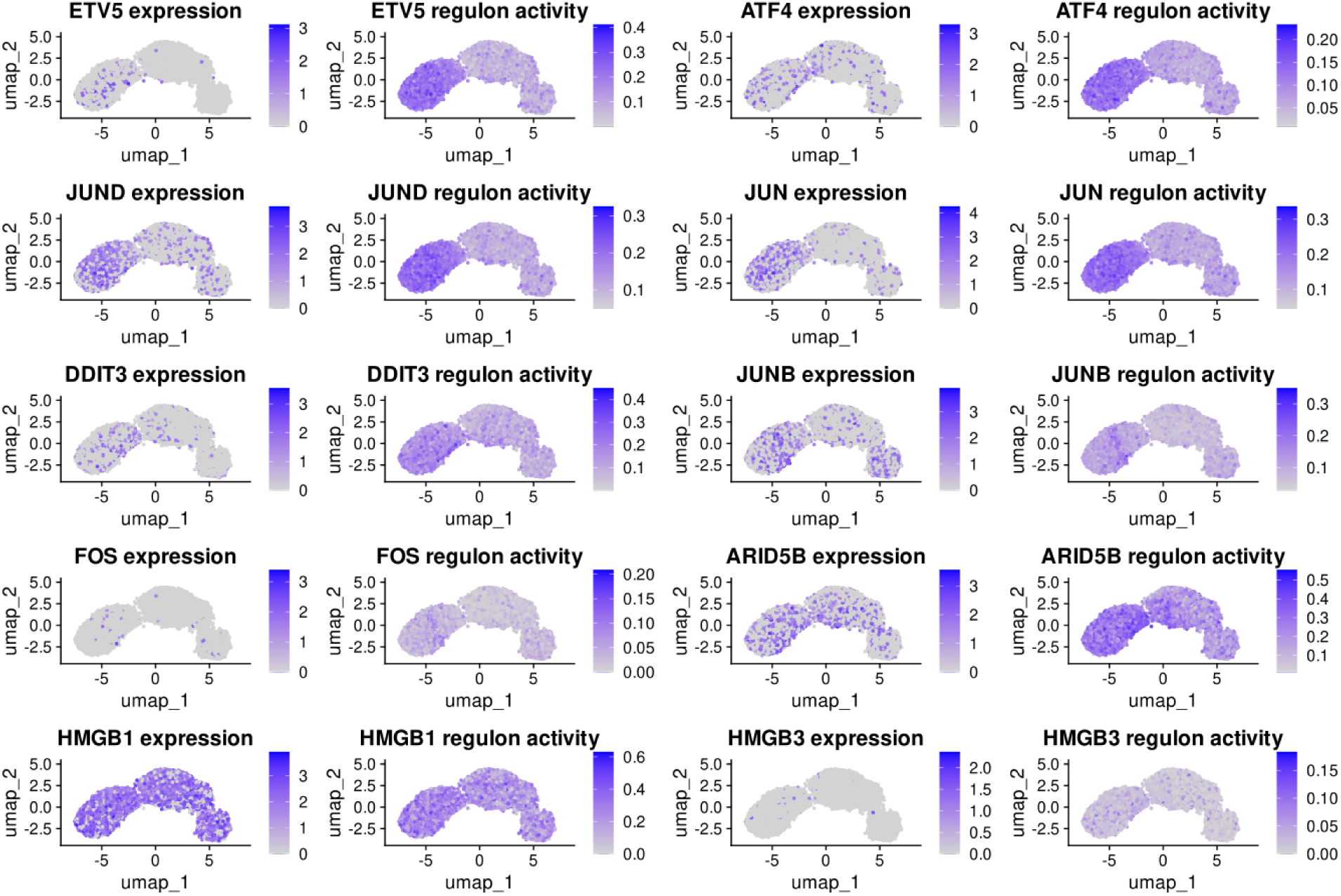
Mapping of the top 10 M27-specific TF expression and regulon activity.

**Figure S3:**
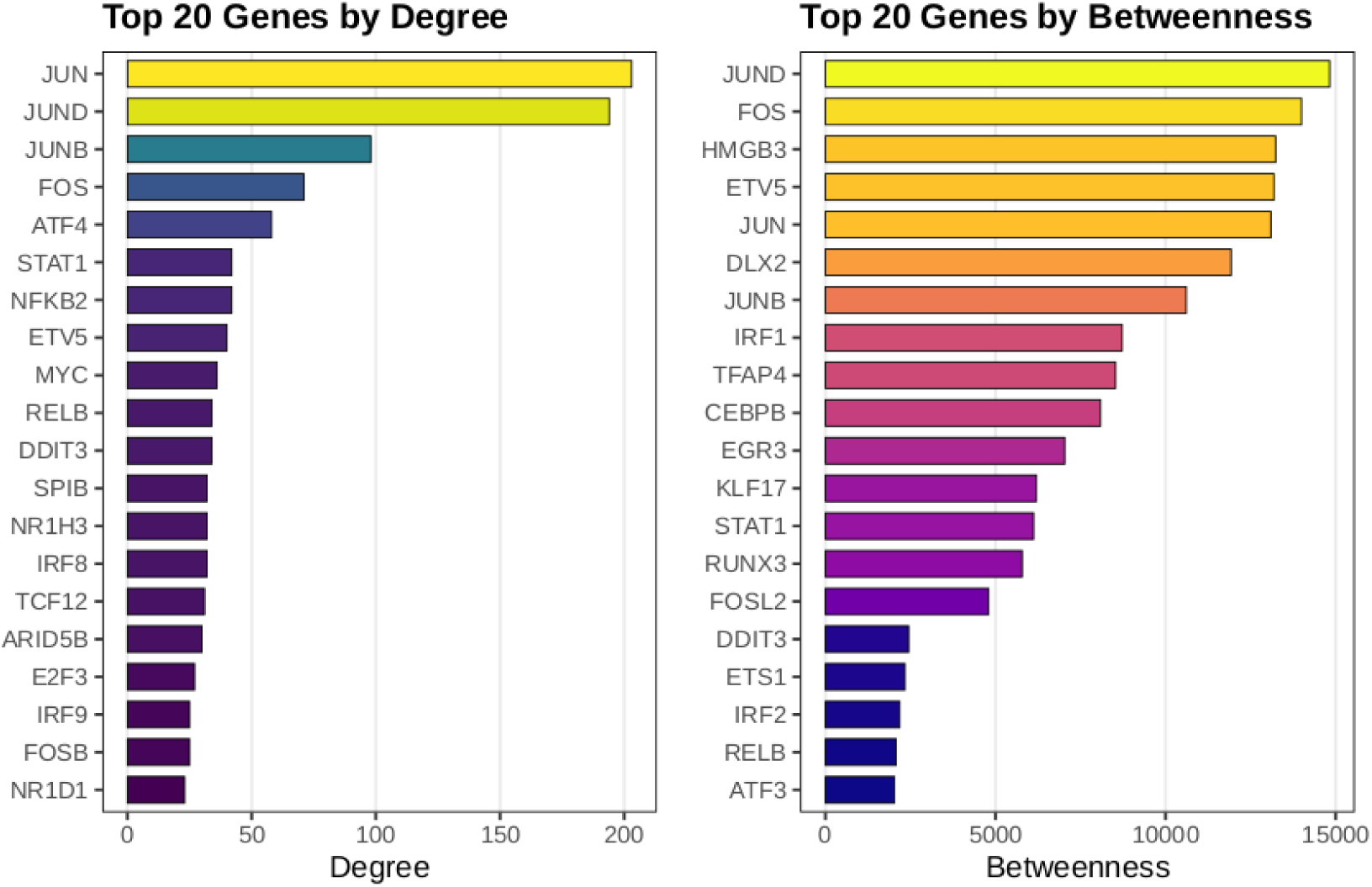
Top 20 genes with the highest degree in the gene regulatory network of CLL cells. Degree (in-degree + out-degree) (left). Betweenness centrality (right).

**Figure S4:**
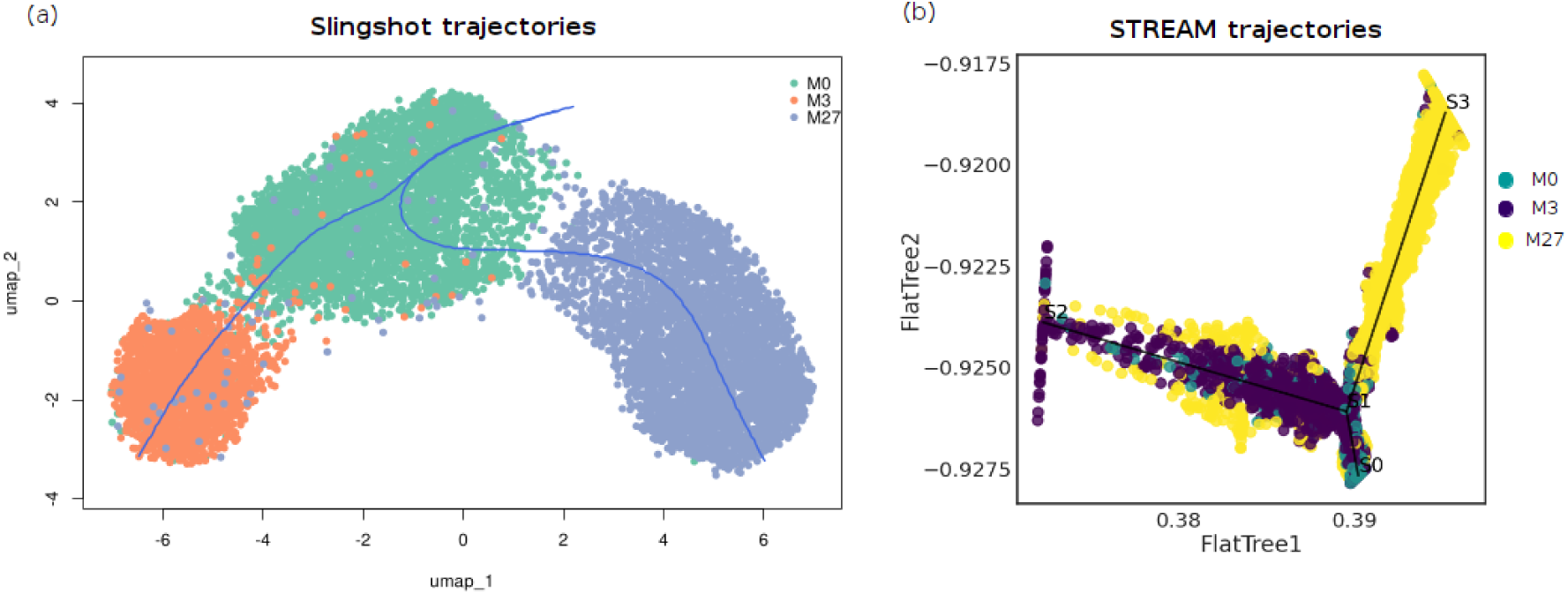
Cell trajectories starting from M0. **(a)** Trajectories inferred with Slingshot. **(b)** Trajectories inferred with STREAM.

**Figure S5:**
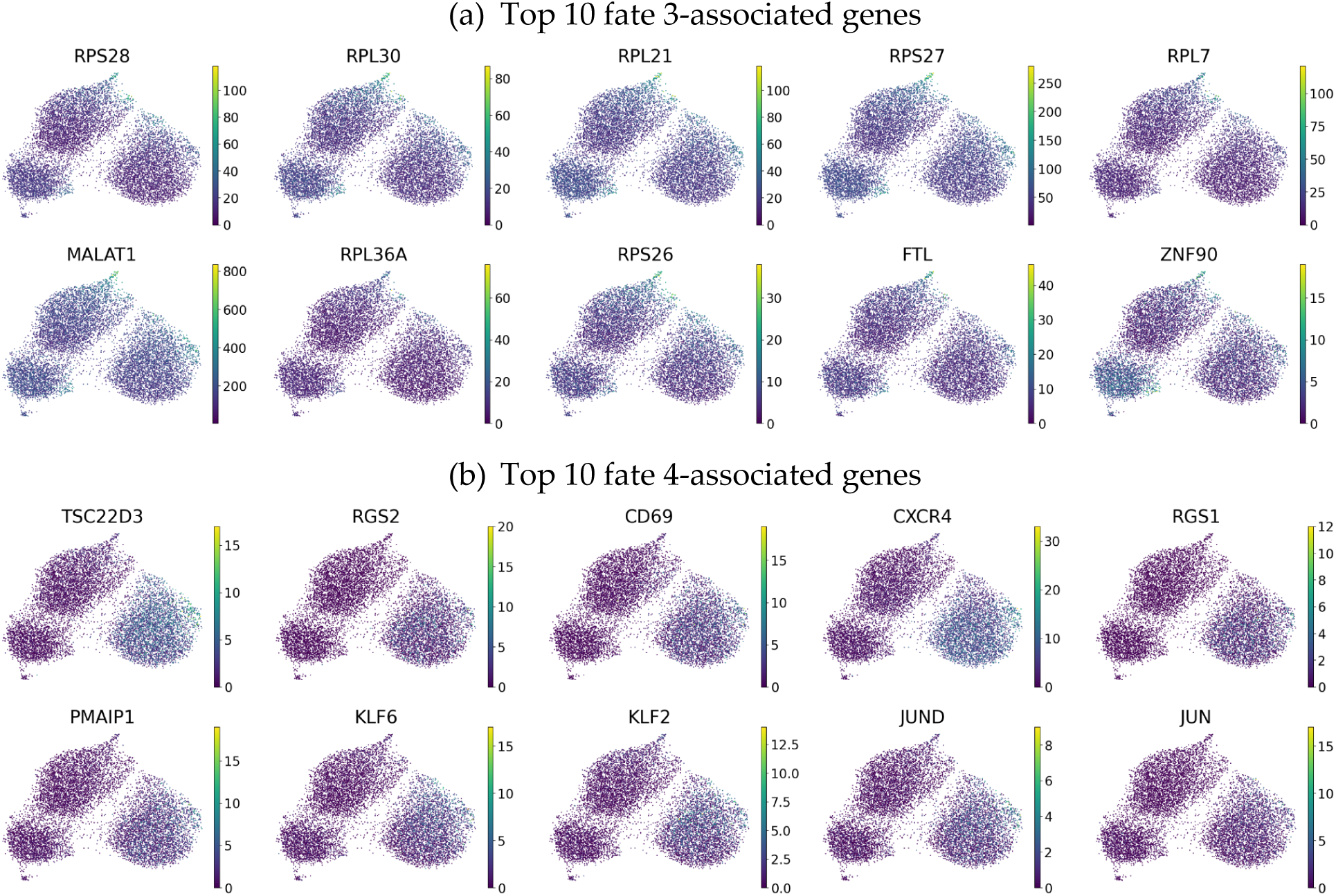
Top 10 (a) response (fate 3) and (b) resistance (fate 4) fate-associated genes.

**Figure S6:**
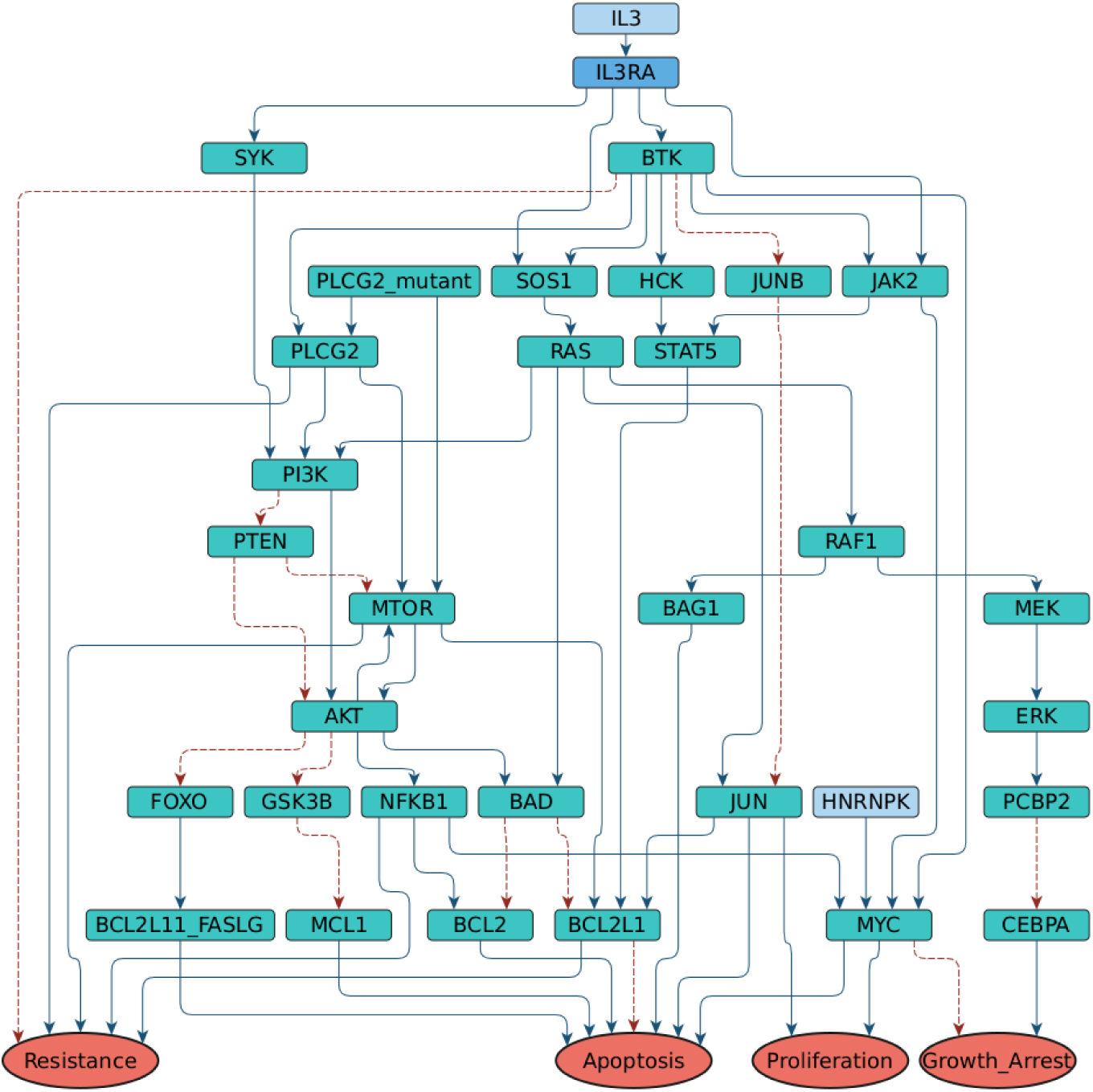
Boolean model of the CLL cells. The model was adapted from (44).

**Figure S7:**
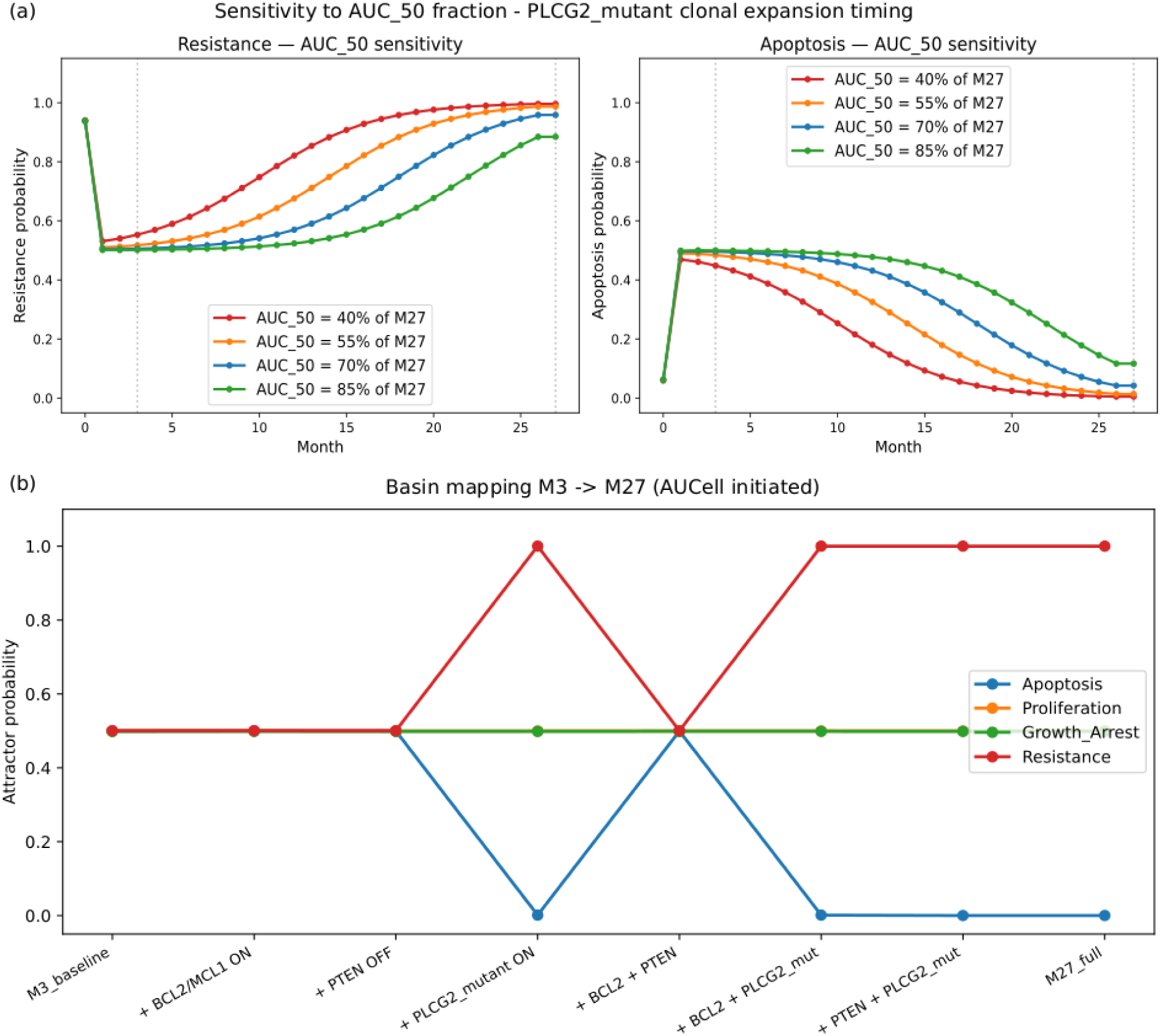
Additional results from the PK-Boolean model.

## Supplementary Material

## SM1. Boolean network construction and pharmacokinetic coupling

### Boolean model construction

#### Starting model and CLL adaptation

We constructed a Boolean network model of CLL B-cell signalling under ibrutinib treatment by adapting the Bio Model Analyser (BMA) leukaemia signalling network (Chuang et al., 2015), originally developed for chronic myeloid leukaemia (CML). We converted continuous-valued BMA update functions to Boolean AND/OR/NOT logic operators and mapped all node names to HGNC-approved gene symbols. The resulting topology served as a structural scaffold, with extensive CLL-specific reengineering, as described below. All Boolean transition rates were set to the binary values {0, 1} throughout, implemented in MaBoSS format (Stoll et al., 2017) and executed via the CoLoMoTo Docker environment (Naldi et al., 2018).

### 1.2 CLL-specific topology modifications

Five categories of modifications were applied to adapt the CML topology to CLL biology (Table 1).

**Table 1.** Summary of major modifications from BMA CML topology to CLL model.

| Category | Nodes affected | Change | Biological rationale |
| --- | --- | --- | --- |
| Oncogenic driver | BCR-ABL1 | Removed; BTK added | BCR-ABL1 absent in CLL; BTK is the central BCR kinase and ibrutinib target (Hendriks et al., 2014) |
| BCR signalosome | SYK, PLCG2, PLCG2_mutant | Added | SYK→PI3Kδ BCR-proximal axis (Young et al., 2019); PLCG2 mutation is a primary ibrutinib resistance mechanism (Liu et al., 2015) |
| Transcription | MYC | Rule rewritten | BTK→NF-κB→MYC (Ezell et al., 2014) and JAK2→MYC (Xie et al., 2002) arms; CML SMAD/IL-3 arm removed |
| Survival | BCL2L1, Resistance | Rules extended | JUN/mTOR→BCL-XL axes; Three-arm Resistance rule (bypass, mTOR, BCL2L1/NF-κB) |
| Dead ends | FLI1, PAG1, STAT3, IL6R, IL6, BCR_ABL1, ABI1_ABI2 | Removed | No path to any output node; Biologically inactive in CLL context |
| Non-CLL nodes | EPOR, GHR, KDR, SRC, CRKL, CBL, SMAD, Wnt subnetwork | Removed | Absent or not relevant in mature CLL B cells; Redundant with SYK-centred topology |

1. The CML-specific oncogenic driver *BCR*-*ABL1* was removed and replaced by *BTK* as the central *BCR* signalling kinase, whose covalent inhibition by ibrutinib constitutes the primary mechanism of drug action (Woyach et al., 2014).
2. SYK was added as the BCR-proximal kinase (IL3RA→SYK), encoding the SYK→PI3Kδ axis that is the relevant upstream activator of PI3K in CLL B cells and the target of fostamatinib (Herman et al., 2010). The PI3K activation rule was updated to reflect the SYK→PI3Kδ and PLCG2→PI3K BCR-signalosome architecture, replacing the CML IL-3RA-dependent rule. The PLCG2 and PLCG2_mutant nodes were added to encode wild-type BCR bypass signalling and the gain-of-function resistance mutation, respectively, consistent with the predominant ibrutinib resistance mechanism in CLL (Woyach et al., 2014).
3. The MYC transcriptional rule was rewritten to encode the BTK→NF-κB→MYC and JAK2/HNRNPK→MYC axes, replacing the CML-derived SMAD/IL-3 combinatorial gate (Herman et al., 2010).
4. BCL2L1 was extended with JUN/ATF2 and mTOR activation arms consistent with documented AP-1→BCL-XL and mTOR→BCL-XL resistance mechanisms (Klanova et al., 2016).
5. The Resistance output node encodes three mechanistically distinct arms: PLCG2-mediated BTK bypass (PLCG2 & !BTK), mTOR-driven survival (MTOR), and NF-κB/BCL-XL co-activation (BCL2L1 & NFKB1).

### 1.3 Iterative model refinement and dead-end removal

The model was refined through two successive cleaning steps. Seven dead-end nodes with no path to any phenotypic output node were removed: FLI1, PAG1, STAT3, IL6R, IL6, BCR_ABL1, and ABI1_ABI2. Two rule corrections were applied: the SMAD→MYC activation edge was restored from the original BMA model, and the Apoptosis→Resistance inhibitory feedback was removed to enforce the design principle that output nodes must not feed back into intracellular signalling rules. A further 15 nodes were removed on CLL-specific biological grounds: EPOR and GHR (cytokine receptors absent on mature B cells), SMAD (TGF-β effector from the CML context), KDR and SRC (VEGFR2/SRC→PI3K axis, minor relative to SYK in CLL), CRKL and CBL (CML-specific BCR-ABL1 substrate adaptors redundant in the CLL BCR context), and the complete Wnt sub-network (FZD, AXIN2, CTNNB1, TCF). The ligand nodes VEGFA, EPO, GH1, and Wnt became dead ends following the removal of their downstream receptors and were also eliminated. Eight logic rules were patched to remove references to deleted nodes (Table 1). The final network comprises 37 nodes and 66 directed edges.

### 1.4 Condition-specific initial states and pharmacological constraints

Three simulation conditions represent the CLL treatment time course: (i) M0 (pre-treatment baseline), (ii) M3 (early ibrutinib response, 3 months), and (iii) M27 (acquired resistance, 27 months). All unconstrained nodes were initialised at equal probability (0.5) in the base configuration file, ensuring that attractor probabilities reflect network dynamics and data-derived constraints rather than manual assignments.

Two categories of pharmacological constraints were applied as condition-specific locks: (i) BTK inhibition, encoded as a continuous transition rate derived from the pharmacokinetic model rather than a binary lock, reflecting the incomplete occupancy achieved at therapeutic trough concentrations; (ii) condition-specific biological states, including PLCG2_mutant=ON and BCL2/MCL1=ON at M27 to encode acquired resistance mechanisms (see Pharmacokinetic-Boolean coupling). Cytokine input (IL3) was treated as active throughout the simulation, consistent with partial microenvironmental cytokine support of circulating CLL cells. Only two nodes were unconditionally excluded from data-driven initialisation: BTK (activity determined by PK) and PLCG2_mutant (somatic mutation state). All other nodes, including the BCL2 family, were initialised at random initial values (*prob*([*node*] = 1) = *prob*([*node*] = 0) = 0. 5).

### 1.5 Model initialisation from AUCell results

To ground the Boolean model TF nodes in single-cell transcriptomic data, regulon enrichment scores derived from AUCell analysis (Aibar et al., 2017) were used to initialise transcription factor (TF) node states. Per-cell AUCell scores were computed for each regulon, binarised per regulon using an automatically selected threshold (valley of bimodal distribution if Sarle’s bimodality coefficient BC > 0.555, otherwise a discriminability-maximising threshold), and then aggregated to a majority-vote consensus per timepoint: a node was assigned binary state 1 if the regulon was active in the majority of cells (≥50%) at that timepoint. The intersection of GRN-inferred TFs and Boolean model nodes was computed at runtime from the .bnd file and the AUCell output, with manual aliases applied where AUCell TF names differed from HGNC node names (e.g. STAT5A/STAT5B → STAT5). The resulting per-condition binary TF states were applied via set_istate() calls, which take lower priority than mutate() pharmacological locks, ensuring that data-driven initialisation never overrides drug mechanism constraints.

### Stochastic simulation framework

#### 2.1 MaBoSS simulation parameters

All simulations were performed using MaBoSS (Markovian Boolean Stochastic Simulator; (Stoll et al., 2017)) via the pyMaBoSS Python API within the CoLoMoTo Docker environment. Each simulation used 100,000 Monte Carlo trajectories (sample_count = 100000), a maximum simulation time of 20 arbitrary units with a time step of 0.1, and 4 parallel threads. Phenotypic output nodes (*Apoptosis*, *Proliferation*, *Growth_Arrest*, *Resistance*) were designated as non-internal; all other nodes were tracked as internal states. Final attractor probabilities were recorded at t = 20 a.u., by which point all simulations had reached stable steady-state distributions.

#### 2.2 Perturbation and combination screens

Single-node perturbation screens were performed exhaustively over all 33 screen-eligible nodes (excluding constitutive inputs, pharmacological lock targets, and output nodes) in both ON and OFF directions. Effect sizes were computed as Δ*Resistance* and Δ*Apoptosis* relative to the corresponding AUCell-initialised reference attractor. Pairwise combination screens used a pathway-stratified design: one node per class (*MAPK*/*ERK*, *PI3K*/*mTOR*, *BCR* bypass, *BCL2* family, *NF-κB*/*AP-1*) was selected as the strongest single-node resistance suppressor, ensuring mechanistic independence between paired nodes. Bliss independence synergy was computed as *B* = *E^AB^* − (*E^A^* + *E^B^* − *E^A^* · *E^B^*), where positive E encodes resistance suppression (*E* =− Δ*Resistance*). Basin of attraction mapping introduced the three major M27 resistance events (*BCL2*/*MCL1* upregulation, *PTEN* loss, *PLCG2_mutant*) individually and in all combinations onto the M3 background. *BTK* inhibition dose-response swept *BTK rate_up* from 0 to 1.0 in 11 steps, with *BCL2* family nodes released to allow a dynamic *Apoptosis* response.

### Pharmacokinetic-Boolean coupling

#### 3.1 PK data and concentration extraction

Individual-patient ibrutinib pharmacokinetic data were obtained from a population PK model providing hourly predicted plasma concentrations (C^pred^, ng/mL) from M0 to M27, combining observed measurements (M0-M6) and model-predicted values (M7-M27). Monthly trough concentrations were defined as the mean of the bottom 10th percentile of hourly *C^pred^* values within each calendar month. The trough was selected as the pharmacologically relevant PK metric for a covalent irreversible inhibitor: at trough, newly synthesised *BTK* protein has accumulated during the dosing interval and represents the pool of unoccupied (pharmacologically active) *BTK*. Cumulative AUC was computed by trapezoidal integration of *C^pred^* from day 0 through each month-end, yielding a monotonically increasing exposure metric used in the clonal expansion model.

### 3.2 PK to PD conversion: BTK inhibition rate

Monthly trough concentrations were converted to BTK Boolean transition rates via a Hill occupancy model:

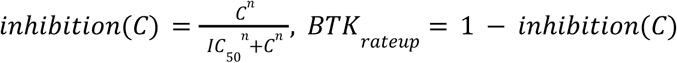

with *IC*_50_ = 1. 0 *ng*/*mL* and Hill coefficient *n* = 1, appropriate for a covalent inhibitor where the kinetic *IC*_50_ for irreversible bond formation approximates a first-order occupancy model (47,48). In MaBoSS, rate_up is the propensity for *BTK* to transition from OFF to ON per unit time; with rate_down fixed at 1.0, the steady-state fraction of trajectories with BTK active equals *rate_up_* /(1 + *rate_up_*). At the observed steady-state trough of ∼1.8 ng/mL, *BTK rate_up* ≈ 0.36, corresponding to ∼26% residual BTK activity, encoding partial rather than complete pharmacological inhibition. Sensitivity of the resistance trajectory to *IC*_50_ was assessed over the range 0.5-5.0 ng/mL.

### 3.3 PLCG2_mutant clonal expansion model

The probability of PLCG2_mutant clonal dominance at month t was modelled as a logistic function of cumulative ibrutinib exposure, reflecting the hypothesis that sustained drug pressure selects for pre-existing PLCG2-mutant subclones:

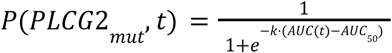

where *AUC*(*t*) is the cumulative trapezoidal AUC from day 0 to the end of month *t* (ng/mL·day), *AUC*_50_ = *f* · *AUC*(*t* = 27) is the cumulative exposure at which *P* = 0. 5, *f* = 0. 7 (calibrated so that clonal dominance coincides with the observed clinical resistance timepoint), and *k* = 8/*AUC*(27) scales the sigmoid to span the full 0→1 range over the observation window. Sensitivity to *AUC*_50_ was assessed over f ∈ {0.40, 0.55, 0.70, 0.85}. At each monthly simulation step, *PLCG2_mutant* was initialised as a Bernoulli random variable with probability *P*(*PLCG*2*_mut_*, *t*), encoding the stochastic representation of mixed wild-type and mutant cell populations within the tumour.

#### 3.4 State-chained monthly simulations

The PK-coupled simulation was structured as a chain of monthly MaBoSS runs from M0 to M27 (28 steps × 50 stochastic sub-steps = 1,400 total simulations). At each step: (i) BTK *rate_up* was set from the monthly trough via the Hill equation; (ii) *PLCG2_mutant* was initialised from *P*(*PLCG*2*_mut_*, *t*); (iii) non-pharmacologically locked nodes were initialised from the mean attractor probability distribution of the previous month (state chaining). State chaining propagates the history of the cell population forward in time: a population that spent previous months in a partially resistant state is more likely to converge to the resistance attractor in subsequent months. At month 0, TF nodes were initialised from AUCell-derived M0 binary states rather than from a prior simulation.

Three parallel trajectory analyses were performed using AUCell-derived starting states for M0, M3, and M27, respectively. The M3 trajectory began at month 3, seeding the first step with AUCell M3 TF binary states combined with M3 pharmacological constraints (*BTK rate_up* from M3 trough, *IL3* active). The M27 trajectory provided a single reference point. This design tests whether cells already in the M3 transcriptional state, or the M27 resistance state, follow the same trajectory as cells starting from M0, providing mechanistic evidence for or against the pre-existing resistance hypothesis.

#### 3.5 Sensitivity analyses

Two sensitivity analyses were conducted. First, the *IC*_50_ parameter was varied over the clinically reported range (0.5, 1.0, 2.0, 5.0 ng/mL) to assess whether the resistance emergence timing depends on assumptions about BTK occupancy. Second, the AUC_50_ fraction f was varied over {0.40, 0.55, 0.70, 0.85} to characterise the sensitivity of the M27 phenotypic transition to the assumed timing of *PLCG2_mutant* clonal dominance. Both analyses used 10 stochastic replicates per condition and the same state-chained monthly structure as the main simulation.

### SM2. Libraries and packages

| Environment |  | Version |  |
| --- | --- | --- | --- |
| R |  | 4.6.1 |  |
| Python |  | 3.13.2 |  |
| Package | Language | Version | Used for |
| Seurat | R | 5.5.0 | Single-cell object handling, preprocessing, DE (FindMarkers) |
| pySCENIC | Python | 0.12.1 | GRN inference pipeline |
| arboreto (GRNBoost2) | Python | 0.1.6 | TF-target regulatory link inference |
| CellRank | Python | 2.0.7 | Trajectory/fate probability inference |
| Slingshot | R | 2.20.0 | Trajectory cross-validation |
| STREAM | Python | 1.1 | Trajectory cross-validation |
| cluster | R | 2.1.8.2 | Silhouette, gap statistic |
| CoLoMoTo Docker | Python | 9.3 | Computational environment for Boolean model execution |
| h2o | R | 3.44.0.3 | AutoML regression: PK features → M0-residualised transcriptomic change |

